# Partial exogastrulation due to apical-basal polarity of F-actin distribution disruption in sea urchin embryo by omeprazole

**DOI:** 10.1101/2021.08.31.458310

**Authors:** Kaichi Watanabe, Yuhei Yasui, Yuta Kurose, Masashi Fujii, Takashi Yamamoto, Naoaki Sakamoto, Akinori Awazu

**Affiliations:** Graduate School of Integrated Sciences for Life, Hiroshima University, Higashi-Hiroshima, 739-8526, Japan; Department of Mathematical and Life Sciences, Graduate School of Science, Hiroshima University, Higashi-Hiroshima, 739-8526, Japan; Research Center for the Mathematics on Chromatin Live Dynamics, Hiroshima University, Higashi-Hiroshima, Hiroshima, 739-8526, Japan

**Keywords:** Gastrulation, partial exogastrulation, sea urchin embryo, actin, omeprazole

## Abstract

Gastrulation is a universal process in the morphogenesis of many animal embryos. Although morphological and molecular events in gastrulation have been well studied, the mechanical driving forces and underlying regulatory mechanisms are not fully understood. Here, we investigated the gastrulation of embryos of a sea urchin, *Hemicentrotus pulcherrimus*, which involves the invagination of a single-layered vegetal plate into the blastocoel. We observed that omeprazole, a proton pump inhibitor capable of perturbing the left-right asymmetry of sea urchin embryo, induced “partial exogastrulation” where the secondary invagination proceeds outward. During early gastrulation, intracellular apical-basal polarity of F-actin distribution in vegetal half were higher than those in animal half, while omeprazole treatment disturbed the apical-basal polarity of F-actin distribution in vegetal half. Furthermore, gastrulation stopped and even partial exogastrulation did not occur when F-actin polymerization or degradation in whole embryo was partially inhibited via *RhoA* or *YAP1* knockout. A mathematical model of the early gastrulation reproduced the shapes of both normal and exogastrulating embryos using cell-dependent cytoskeletal features based on F-actin. Additionally, such cell position-dependent intracellular F-actin distributions might be regulated by intracellular pH distributions. Therefore, apical-basal polarity of F-actin distribution disrupted by omeprazole may induce the partial exogastrulation via anomalous secondary invagination.

## 1 Introduction

Gastrulation is an essential morphogenic process in various animals, wherein a ball of single-layered cells (blastula) differentiates into a multilayered gastrula or early embryo (Stower and Bertocchini, 2017; Shindo, 2018; Nájera and Weijer, 2020). Species-specific variations in this process provide the basis for particular animal morphologies and lead to the generation of internal organs. The basic mechanism of whole embryonic transformations induced by organizational rearrangement and cell movements are evolutionarily conserved.

Sea urchin embryos are commonly used for studying various morphogenetic processes universally observed in animals, such as the left-right symmetry breakage (McCain et al., 1994; Aihara and Amemiya, 2001; Warner and McClay, 2014; Takemoto et al., 2016), neural network formations (Yaguchi et al., 2010; Burke et al., 2014; McClay et al., 2018), and gastrulation (Dan and Okazaki, 1956; Gustafson and Kinnander, 1960; Hardin and Cheng, 1986; Kominami and Takata, 2004; Martik and McClay, 2017) due to its evolutionary position as diverged at early period of deuterostome evolution. Furthermore, the gene regulatory network controlling endomesoderm specification in sea urchin embryos have been well studied (Davidson et al., 2002; Oliveri and Davidson, 2004). Sea urchin gastrulation progresses through the following five steps (Kominami and Takata, 2004): Step 1 (after hatching), the embryo becomes an epithelial monolayer with a thickened vegetal plate; step 2 (primary invagination), the vegetal plate bends to invade the blastocoel and a short tubular archenteron is formed; step 3 (lag phase in archenteron elongation), secondary mesenchyme cells (SMCs) appear at the tip; step 4 (secondary invagination), the archenteron elongates by pulling on the filopodia of the SMC and cell rearrangement; step 5 (tertiary invagination), presumptive endodermal cells are recruited into the archenteron. Steps 1 and 2 have been conventionally referred to the primary invagination, and steps 3 and 4 to the secondary invagination (Dan and Okazaki, 1956; Gustafson and Kinnander, 1956).

Embryos of some sea urchin species exhibit anomalous morphogenesis (exogastrulation) under various treatments where the vegetal plate of the embryo prolapses outward instead of invaginating. For example, the archenteron of embryos cultured in the presence of LiCl or sugar is completely evaginated due to the depolymerization of SMC microtubules in the blastocoel space connecting the primary intestine and ectoderm (Dan and Okazaki, 1956; Hardin and Cheng, 1986; Khurrum et al., 2004). These studies suggest that lifting the primary intestine from the vegetal plate by microtubules plays an important role in step 4 of gastrulation. Exogastrulation was also observed via Rab35 knockdown, which was also suggested to disturb whole embryonic cytoskeleton distribution (Remsburg et al., 2021).

Moreover, the vegetal plate bending in step 2 of normally cultured embryos of the sea urchin, *Lytechinus pictus*, occurs even when it is surgically isolated (Ettensohn, 1984). It is suggested that the cells at the archenteron tip should be bottle-shaped (Nakajima and Burke, 1996), and those around the vegetal plate should be wedge-shaped (Burke et al., 1991) to initiate primary invagination (Kominamiand Takata, 2004). Additionally, mathematical models of vegetal poles suggest that primary invagination occurs under the appropriate force conditions for “apical constriction,” “cell tractor,” “apical contractile ring,” “apicobasal contraction,” and “gel swelling” (Odell et al., 1981; Davidson et al., 1995). However, it is unclear which of these effects are essential for development after primary invagination (steps 2).

In this study, the roles of the dominant factors in steps 2 and after of sea urchin gastrulation are determined through experimental analysis and mathematical modeling. First, omeprazole that was known as the gastric acid suppressant inhibiting the proton pump activity and as to perturbs left-right asymmetry of sea urchin embryo (Hibino et al., 2006), caused “partial” exogastrulation without the loss of microtubules in the blastocoel space of *Hemicentrotus pulcherrimus* (HP) embryos. Second, the intracellular apical-basal polarity of F-actin distribution in vegetal side cells was stronger than those in animal side cells during the early gastrulation stage (steps 1 and 2), but omeprazole disturbed the polarities in vegetal side. Therefore, the partial exogastrulation was likely due to anomalous cytoskeletal behaviors. Third, CRISPR-Cas9-mediated knockout of cytoskeleton-related genes were performed in HP embryos and the apical-basal ratio of intracellular pH was analyzed using fluorescence imaging. Finally, simulations of morphogenic processes of normal and omeprazole-treated (exhibiting exogastrulation) HP embryos during the pre-early gastrulation were performed using a mathematical model utilizing the cytoskeletal force parameters determined using the above experiments.

## 2 Results

### 2.1 Omeprazole treatment of HP embryos caused partial exogastrulation

The HP embryos exposed to omeprazole from immediately after fertilization and through development exhibited anomalous morphogenesis with partial exogastrulation (Fig. 1). Detailed comparisons of the structural features around the vegetal pole between the normal (control) and treated embryos at each gastrulation step were performed. The following steps were designated based on those defined in a recent report (Kominami and Takata, 2004). In this case, the original step 3 is not shown because the step 3 was obtained at 21-23 h post fertilization (hpf) that overlaps step 2 (20-22 hpf) and step 4 (mid-gastrula: 22-24 hpf).

**Figure 1.**
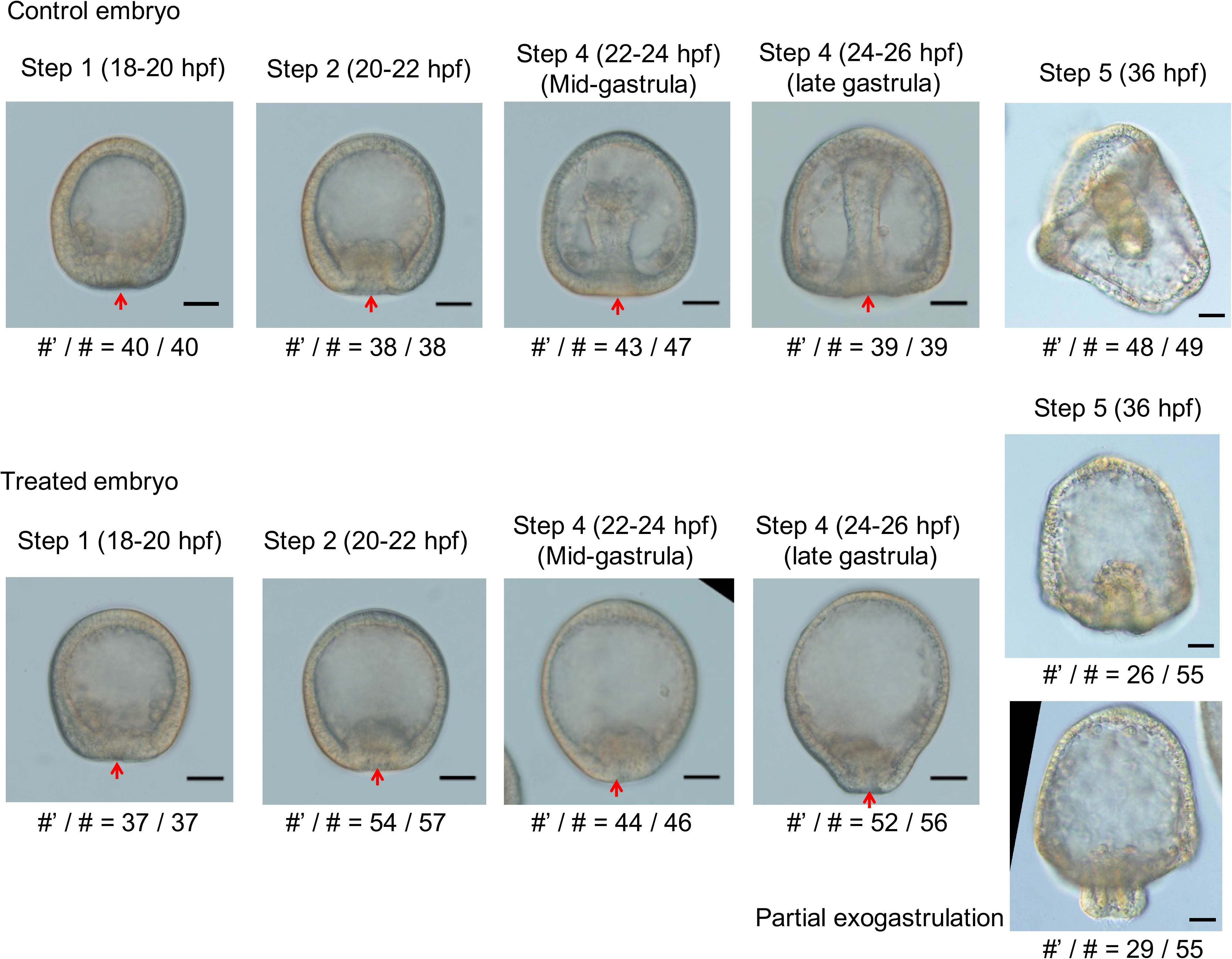
Developmental stages of control and omeprazole-treated HP embryos. Typical bright-field images of the gastrulation process (steps 1, 2, 4, and 5) of sea urchin embryos (scale bars: 30 μm). Arrows indicate vegetal pole positions. Gastrulation did not progress after step 2 in the treated embryos, and the outward protrusion of the vegetal pole side was more pronounced at step 4. In step 5, gastrulation stopped incompletely without penetrating the archenteron, and “partial” exogastrulation was observed in more than half of treated embryos (bellow image). # and #’ refer to the total number of sampled embryos and number of embryos with similar shape to the image, respectively.

In step 1, control and treated embryos exhibited equivalent vegetal pole thickening (18-20 hpf). In step 2, primary invagination occurred in both control and treated embryos at the same time. However, intestinal invagination was slightly shallower in the treated embryos than that in the control embryos (20-22 hpf). In the mid-gastrula stage (step 4; secondary invagination stage), treated embryos did not show further elongation of the archenteron into the blastocoel, and the shape of treated embryos elongated along the animal-vegetal axis (22-24 hpf). Meanwhile, in the control embryos, the archenteron was further elongated, and the presence of SMCs was obvious. In the late gastrula stage (step 4; secondary invagination stage), the outward protrusion of the vegetal pole side was pronounced in the treated embryo (24-26 hpf). The treated embryo had a comparatively little elongation of the archenteron into the blastocoel compared to the control embryos. In step 5 (36 hpf), gastrulation was arrested without penetrating the archenteron in the treated embryos. More than half of treated embryos exhibited “partial” exogastrulation, in which the tip of the archenteron folded inward due to normal primary invagination, but the remaining part extended outward during the secondary invagination. Here, the bottom half of archenteron was stained for endogenous alkaline phosphatase activity in control embryos (Fig. 2a). On the other hand, in treated embryos showing partial exogastrulation, the part extending outward was stained (Fig. 2a), indicating that the part corresponding to endodermal tissue was formed at the outer of these embryos. However, a part of the endodermal tissue was invaginated inward, suggesting that exogastrulated endodermal tissue may be lifted inward in the treated embryos after the secondary invagination.

**Figure 2.**
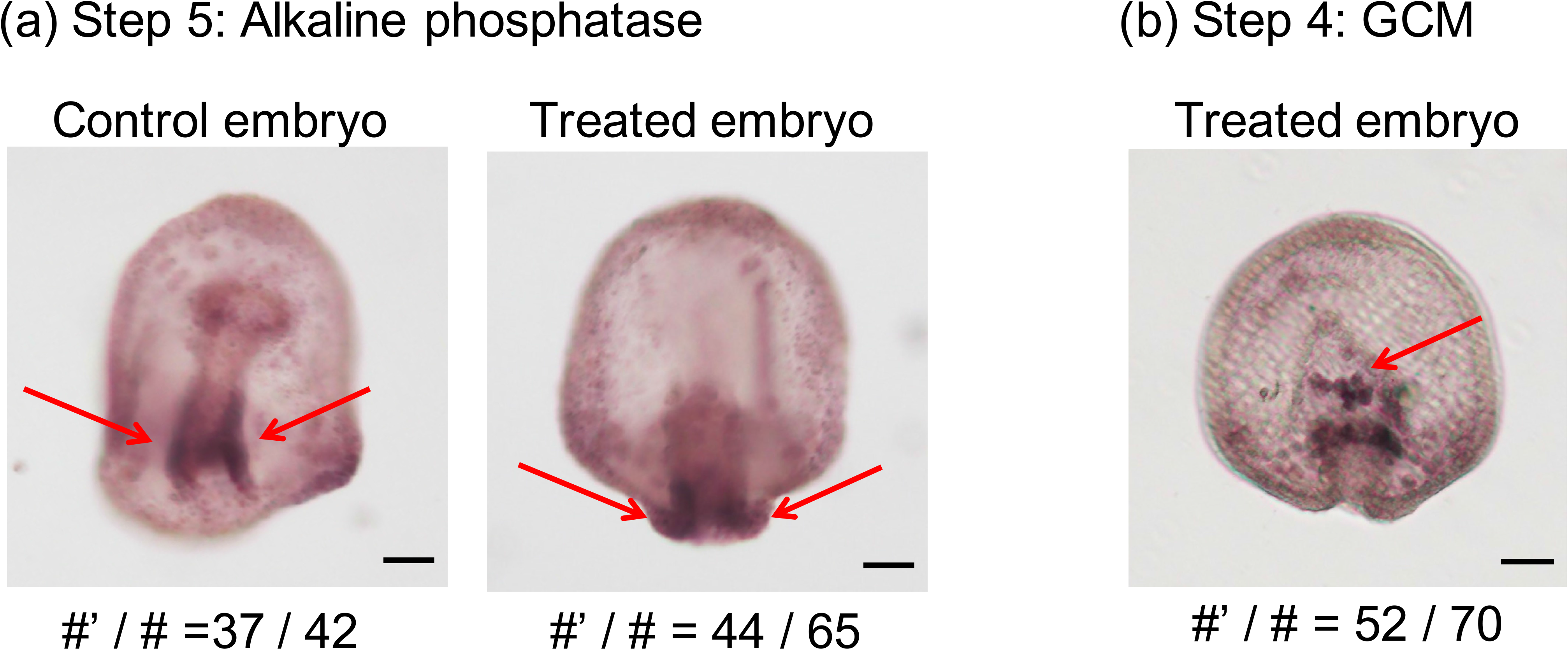
Visualization of endodermal tissue and SMCs. (a) Endodermal tissue was stained for endogenous alkaline phosphatase (red arrows) in control (left) and treated embryos (right) in step 5, where stained part located inner region in control embryos while stained located outer region in treated embryos (scale bars: 30 μm). (b) Typical locations SMCs in treated embryo in step 4 where *gcm*, the marker of SMC, was stained (red arrow) (scale bars: 30 μm). # and #’ refer to the total number of sampled embryos and number of embryos clearly stained similar to the image, respectively.

During the secondary invagination phase, SMCs are detached from the tip of the archenteron and form contractile units; the connection between the archenteron tip and the animal pole tissue via the SMC and pseudopodia facilitates the traction of the archenteron (Dan and Okazaki, 1956; Hardin, 1988). In both control and treated embryos, the migration of SMCs from the tip of archenteron into the blastocoel was observed (Fig. 2b), indicating that the SMCs migrated into the blastocoel of the treated embryo between step 2 and 4 and pseudopodia were formed in the blastocoel.

Previously reported exogastrulation was induced by the loss of pseudopodia (Dan and Okazaki, 1956; Hoshi, 1979; Khurrum et al., 2004); therefore, partial exogastrulation in this report was expected to occur through a different mechanism. On the other hand, whole mount *in situ* hybridization showed the release of SMCs from archenteron in the treated embryos (Fig. 2b). This result suggested that further lift of the archenteron into the blastocoel was induced by SMCs in the treated embryos.

### 2.2 Omeprazole perturbed cytoskeletal distributions in vegetal side cells

The intracellular distribution of F-actin was analyzed at the primary invagination stage (steps 1 and 2) to reveal the mechanism underlying the morphological changes in the treated embryos at the secondary invagination stage (step 4). F-actin was visualized via fluorescence imaging using actinin-GFP where actinin was known as one of F-actin binding proteins (Edlund et al., 2001). The apical-basal polarity of intracellular F-actin distribution was evaluated by estimating the fluorescence intensity ratio between the apical and basal sides of cells (Fig. 3a).

**Figure 3.**
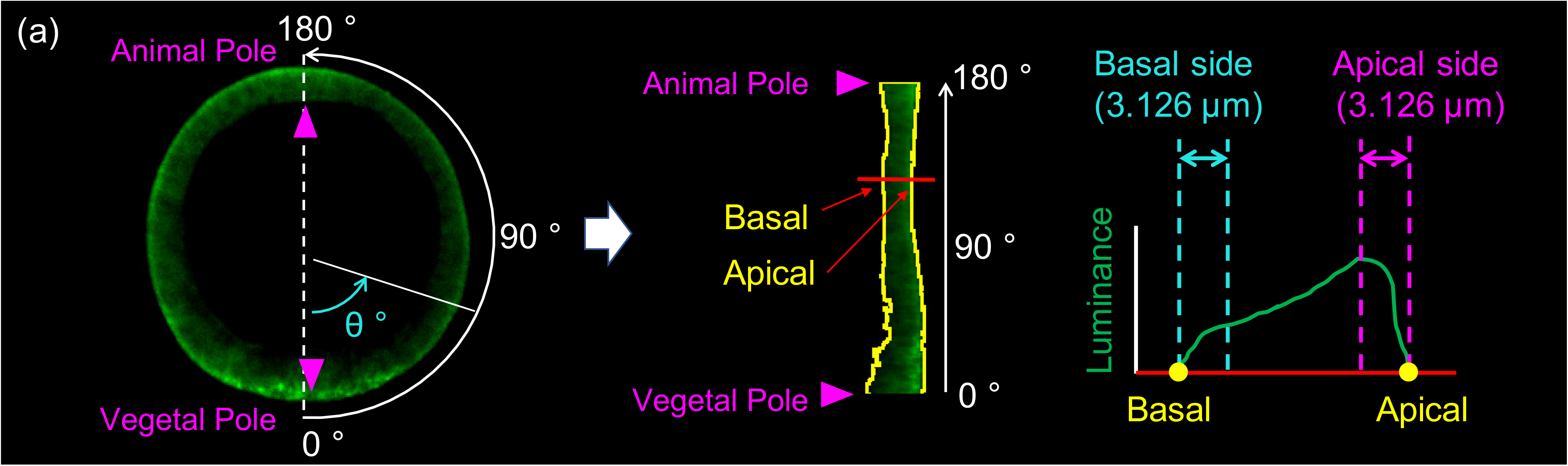
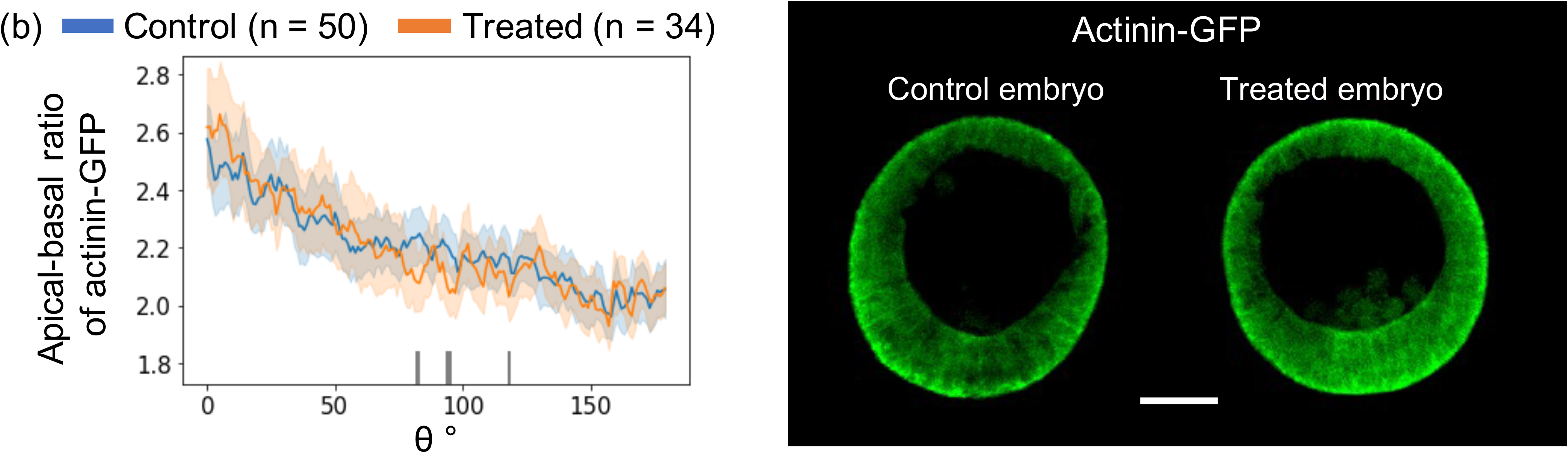
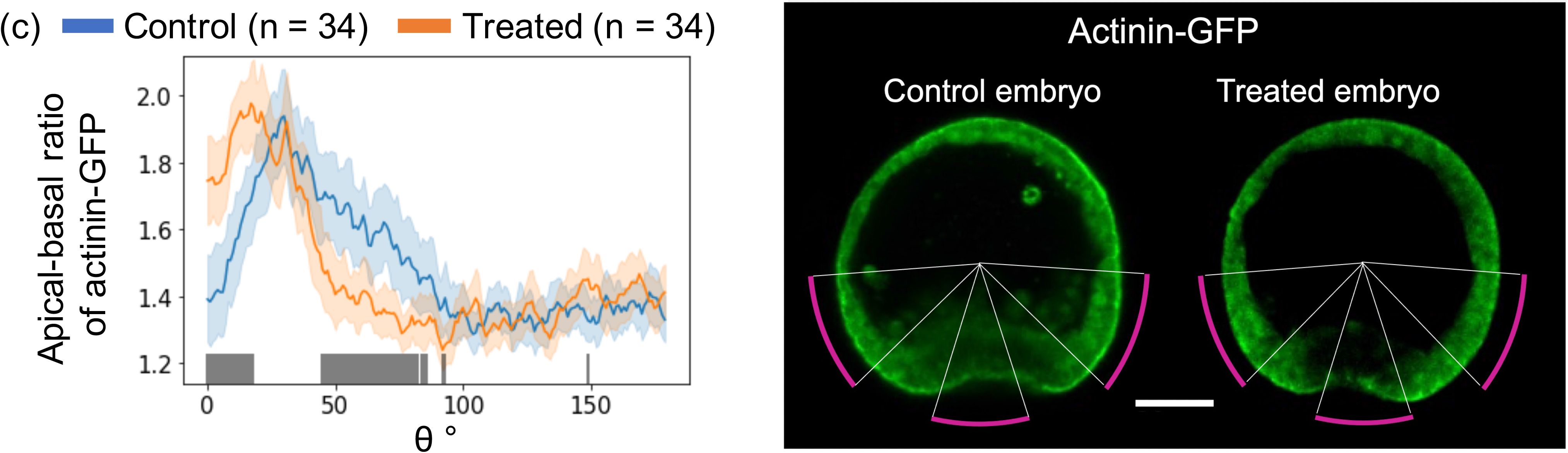
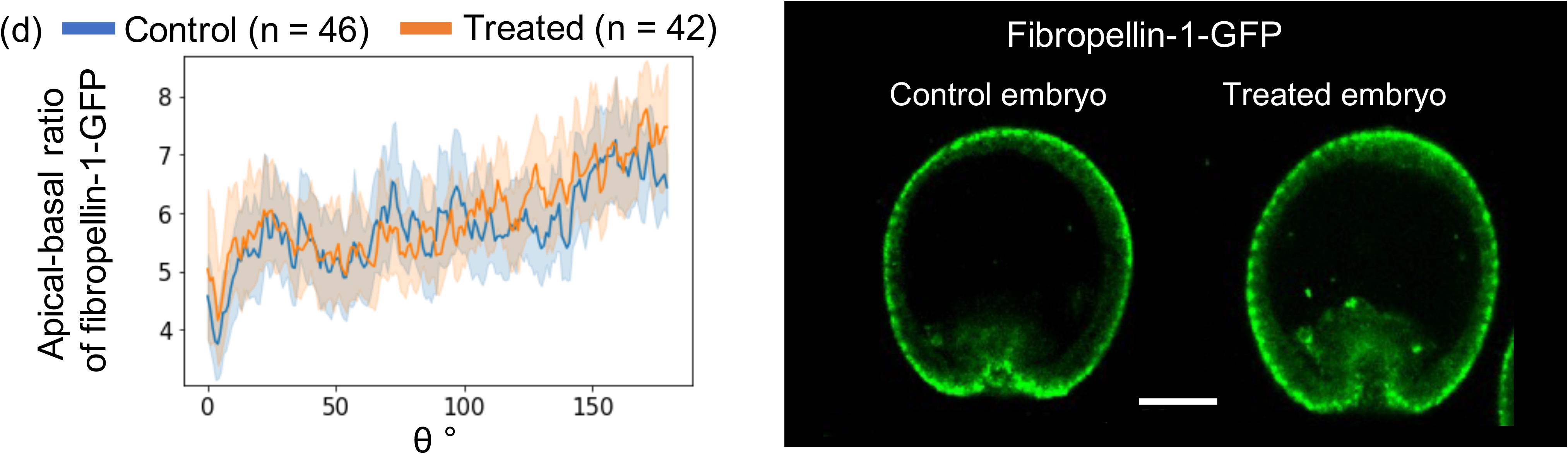
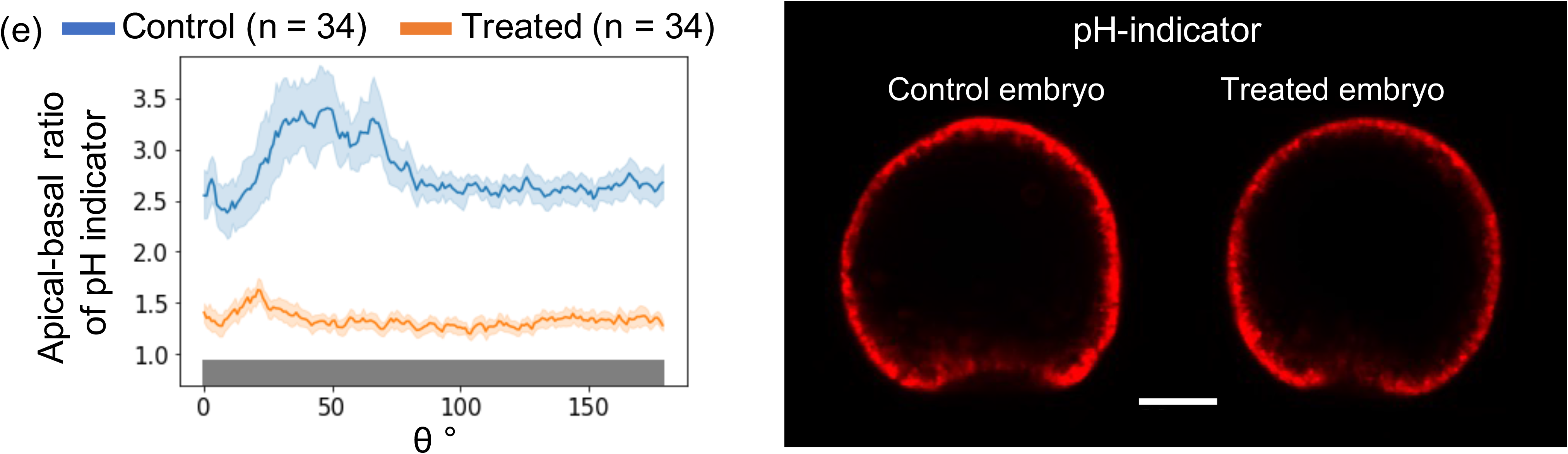
Whole embryonic distributions of actinin-GFP, fibropellin-1-GFP, and pH. (a) Definitions of angle θ (0°-180°) from the vegetal pole (0°) to the animal pole (180°) along the circumference of the embryo cross-section and apical and basal sides of cells (See also Fig. S7) of confocal fluorescence microscopy images determined via actinin (actinin-GFP: green) intensity of embryos. **(b-e)** Average and 95% confidence intervals (error bars) of apical-basal ratios at angle θ obtained by n samples (left), and typical confocal fluorescence microscopy images (scale bars: 30 μm) (right) of actinin-GFP intensities at step 1 **(b)** and step 2 **(c)**, fibropellin-1-GFP intensities at step 2 **(d)**, and pH indicator intensities at step 2 **(e)**. In left figs, blue and orange curves and bars represent the control and treated embryo values, respectively, where gray bars indicate significantly different average values between the control and treated embryos according to Welch’s *t*-test (p < 0.05) (see also Fig. S1). Magenta curves in the right panel of (**c**) were included along the region with significant differences between the values of the control and treated embryos. The correlation coefficients of apical-basal ratios between actinin-GFP intensities **(c)** and pH indicator intensities **(e)** in control and treated embryos were 0.56 and 0.64, respectively.

Similar apical-basal ratio distributions of the actinin-GFP signal were obtained between the control and treated embryos, while the ratios gradually increased closer to the vegetal pole in both embryos at step 1 (Fig. 3b, Fig. S1a, Fig. S2a-d). The distribution of the apical-basal ratios of actinin-GFP signal intensities in step 2 was similar between control and treated embryos only in the animal half (Fig. 3c, Fig. S1b, Fig. S2e-h). However, from the equator to the vegetal pole, the ratio in the control embryos was larger than that in the treated embryos, whereas at the vegetal pole and its surrounding region, control embryos showed a smaller ratio than the treated embryos. Therefore, the formation of an anomalous embryonic shape of treated embryos could correlate with the deviation of apical-basal polarity of F-actin distribution in the vegetal half of the embryos during primary invagination stage. Meanwhile, omeprazole did not influence the whole embryonic distribution of fibropellin-1 (Fig. 3d, Fig. S1c), an F-actin scaffold in the apical lamina of the apical pole in each cell (Nakajima and Burke, 1996; Burke et al., 1998).

Since omeprazole is a proton pump inhibitor, intracellular pH was estimated using a pH indicator; fluorescent intensity increased as pH decreased. In both control and treated embryos in step 2, the apical-basal ratio of the pH indicator was greater than 1 (Fig. 3e, Fig. S1d, S3), indicating that the pH of the apical side of each cell was always lower than that on the basal side. Additionally, this ratio was higher in the vegetal half than that in the animal half of the control embryo, while this pattern disappeared in the treated embryo. Thus, the apical-basal polarity of the pH indicator fluorescent intensity was positively correlated with that of actinin-GFP signal intensity in whole embryos (Fig. 3c, e).

### 2.3 Gastrulation starts but does not progress as normal in cytoskeleton regulator-knockout embryos

CRISPR-Cas9-mediated knockout of the typical enhancer and repressor of F-actin formation, *RhoA* and *YAP1* (Beane et al., 2006; Dupont et al., 2011), were performed by microinjection of *Cas9* mRNA and sgRNAs designed (Fig. S4) to elucidate the contribution of F-actin to gastrulation. Although the mutation frequencies of *RhoA* and *YAP1* knockouts were 55.6 and 77.8%, respectively, and the frameshift rate was 22.3% in either knockout (Fig. S4), gastrulation stopped at step 2 at the primary invagination stage in *RhoA* or *YAP1* knockout embryos (Fig. 4a); further extension of the archenteron was not observed in both knockout embryos. Interestingly, the phenotypes between these knockout embryos were similar, although the gene functions were different (Fig. S4). Pigmented cells were observed at 45 hpf, indicating that development had not ceased. This showed that primary invagination occurred even in embryos with mild perturbation of F-actin function, but subsequent gastrulation processes required appropriate F-actin-derived forces.

**Figure 4.**
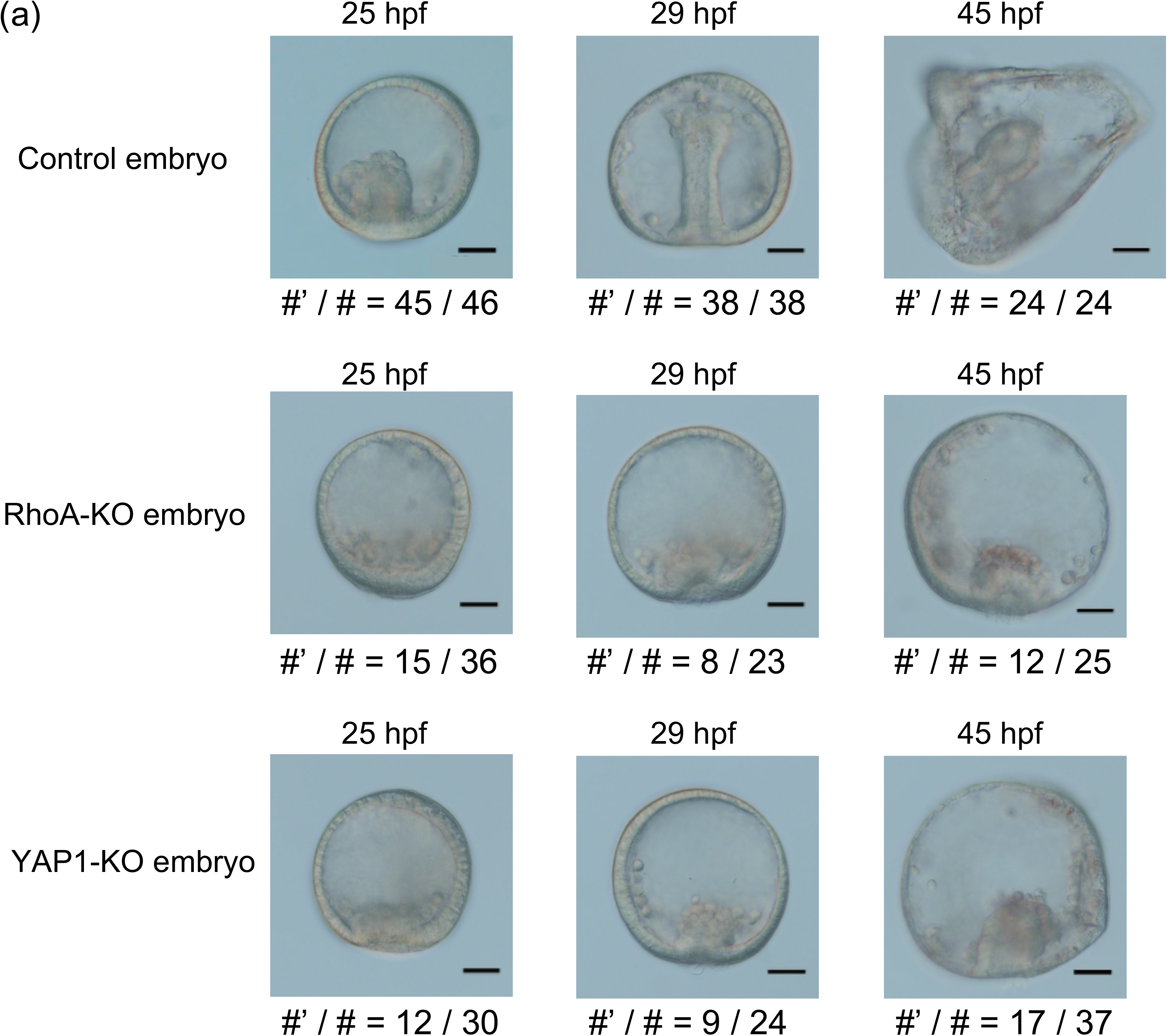
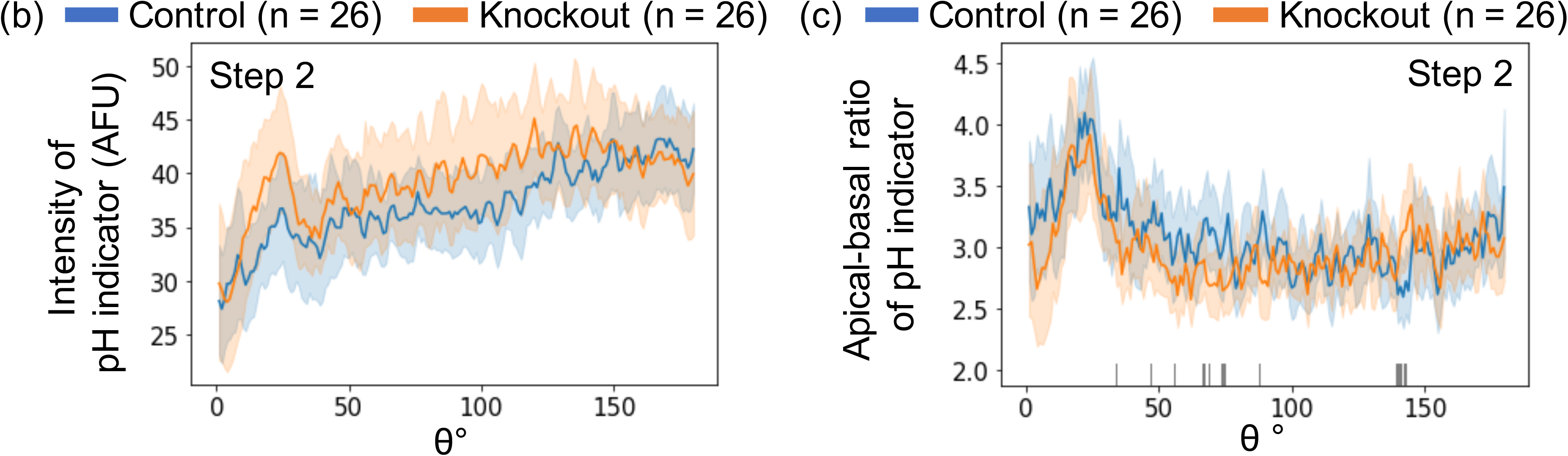
Effect of F-actin regulator-knockout on gastrulation. **(a)** Bright-field images of gastrulation in control, *RhoA*-knockout, and *YAP1*-knockout embryos at selected time-points. PMCs and pigment cells were observed in all embryos suggesting that development did not stop. The knockout embryos did not form the structure like prism larva observed at 45 hpf in the control embryo (scale bars: 30 μm). # and #’ refer to the total number of sampled embryos and number of embryos with similar shape to the image, respectively. **(b-c)** Average fluorescence intensities (arbitrary fluorescence units) and 95% confidence intervals (error bars) of intracellular pH indicator **(b)** and apical-basal ratio of pH indicator **(c)** of *RhoA*-knockout embryos and control embryos as a function of angle θ. The indications of colors and θ are stated in Figures 3 and S1.

The whole embryonic distribution of average intensity and apical-basal ratios of the pH indicator were unchanged in *RhoA*-knockout embryos compared with those in the control, and therefore, perturbation of F-actin polymerization did not affect the pH gradient of the cell (Fig. 4b-c, Fig. S1e-f).

### 2.4 Mathematical model of embryonic shape formations during early gastrulation considering cell-dependent apical-basal intracellular polarity of F-actin distribution

A mathematical model of cell motion at the cross-section including animal and vegetal poles during steps 1-4 of gastrulation was constructed to examine the influence of intracellular F-actin distribution on the formation of embryonic shapes. The model was constructed based on the gastrulation model of *Nematostella vectensis* (Tamulonis et al., 2011) that consisted of cells constructed using springs and beads circularly connected to form a two-dimensional cross-section of the embryo. The following assumptions were made based on experimentally known facts:

I. Each embryo contained three cell types as follows: pigment cells (Kimberly and Hardin, 1998; Kominami and Takata, 2004), wedge cells (Burke et al., 1991), and other cells (Fig. 5). The pigment cells near the cavity entrance were considered to be bottle-shaped (Kimberly and Hardin, 1998; Kominami and Takata, 2004) because of the site-specific force in these cells. This may explain the primary invagination occurring in a surgically isolated vegetal plate (Ettensohn, 1984). In the knockout experiment shown in Fig. 4, partial F-actin function may be enough for the formation of bottle-shaped pigment cells and subsequent primary invagination. In the present model, such effect was represented by the fact that the apical side of the pigment cell was assumed smaller than those of other cells, and the apical side of each wedge cell was assumed larger than the basal side (Fig. 5a). The apical and basal sizes of cells in the initial state (Fig. 5b) were determined by those with which the shape of the model after the relaxation of interaction energy could imitate the embryo shape in step 1 (Fig. 1).
II. The width of the apical and basal sides changed in a cell-dependent manner in control and treated embryos due to cytoskeletal forces generated by F-actin. In the present model, the F-actin concentration in each side of the cell was assumed to correlate positively with the cell cortical force generations that push out each cell side area to stretch tissues due to F-actin polymerization (Pollard et al., 2000, Footer et al., 2007, Mullins et al., 2013). Thus, the lengths of the apical and basal sides of the cells, except pigment cells and wedge cells, were determined by referring the profile of the apical-basal ratio of actinin-GFP in step 2 (Fig. 5a). Here, the elongations of the apical side of these cells were determined based on the apical-basal ratio of actinin-GFP, and the length of the basal side was assumed to change oppositely to the apical side to maintain the cell area.
III. Cell divisions and three-dimensional mutual cell invasions mainly contributing to late gastrulation were not included because these processes were rarely observed in early gastrulation (Mizoguchi, 1999). Therefore, the present model described embryo shape dynamics of the control and treated embryos introducing two-dimensional motions and deformations of the apical and basal sides in 64 cells (Fig. 5a).

**Figure 5.**
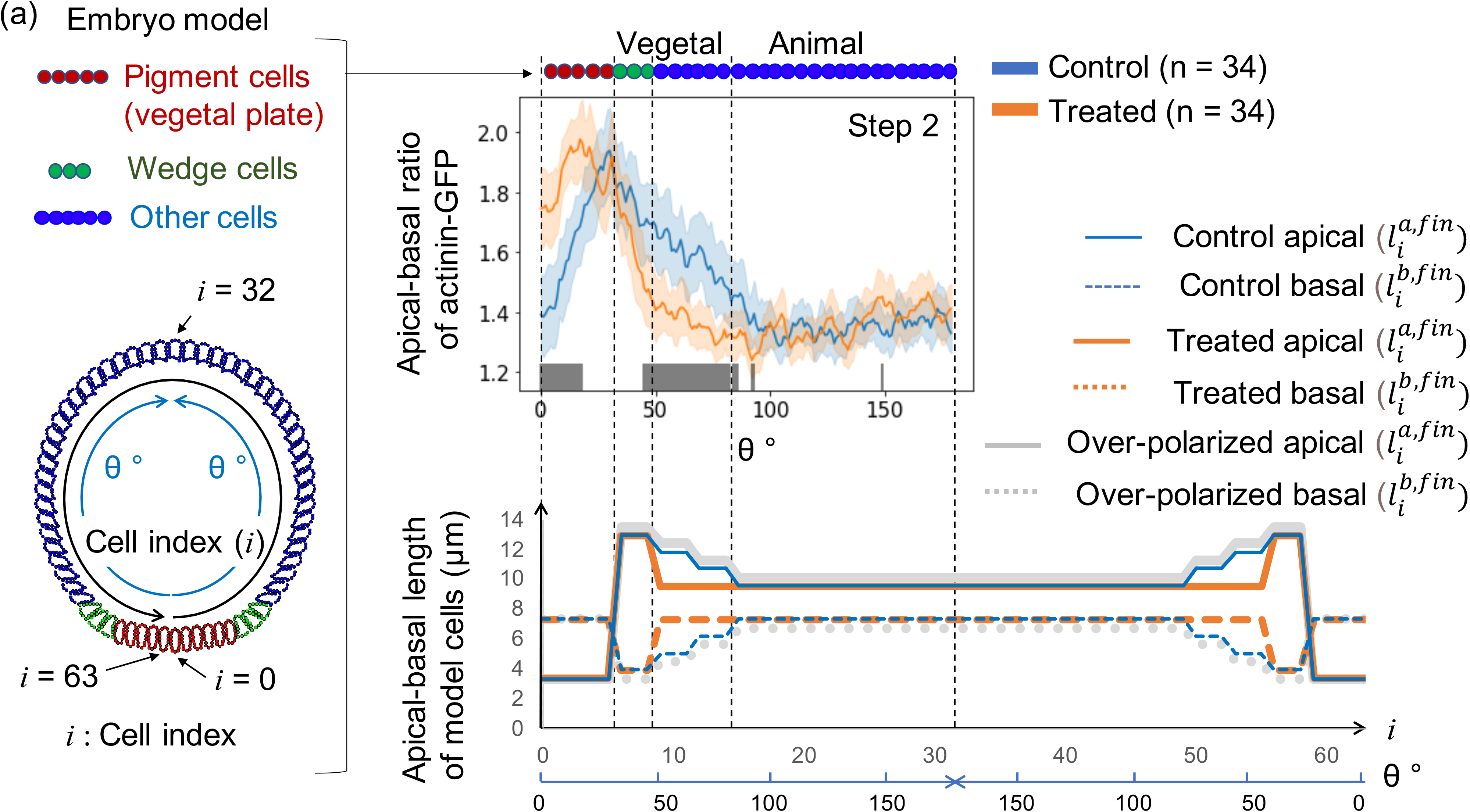
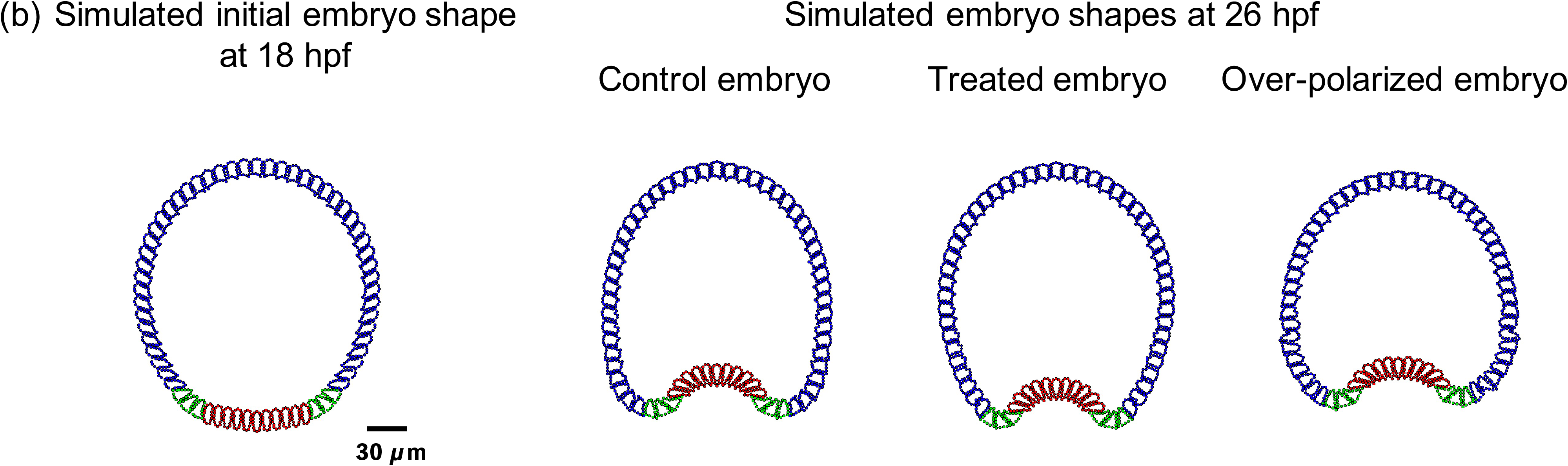
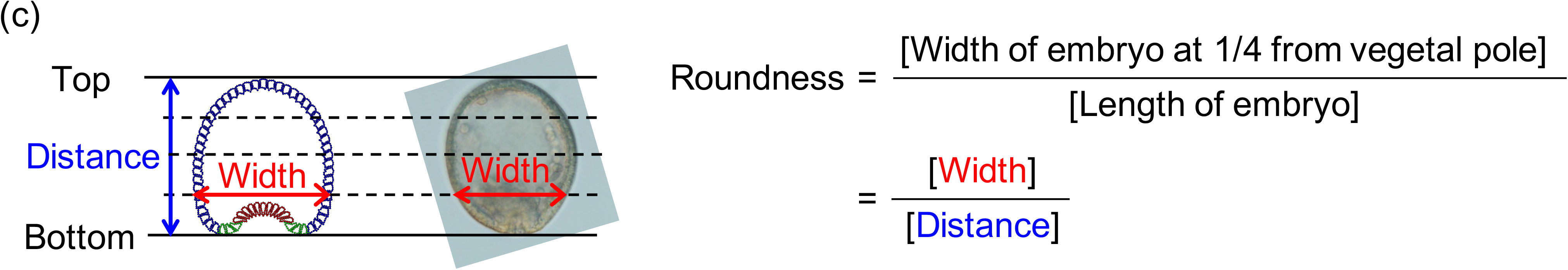
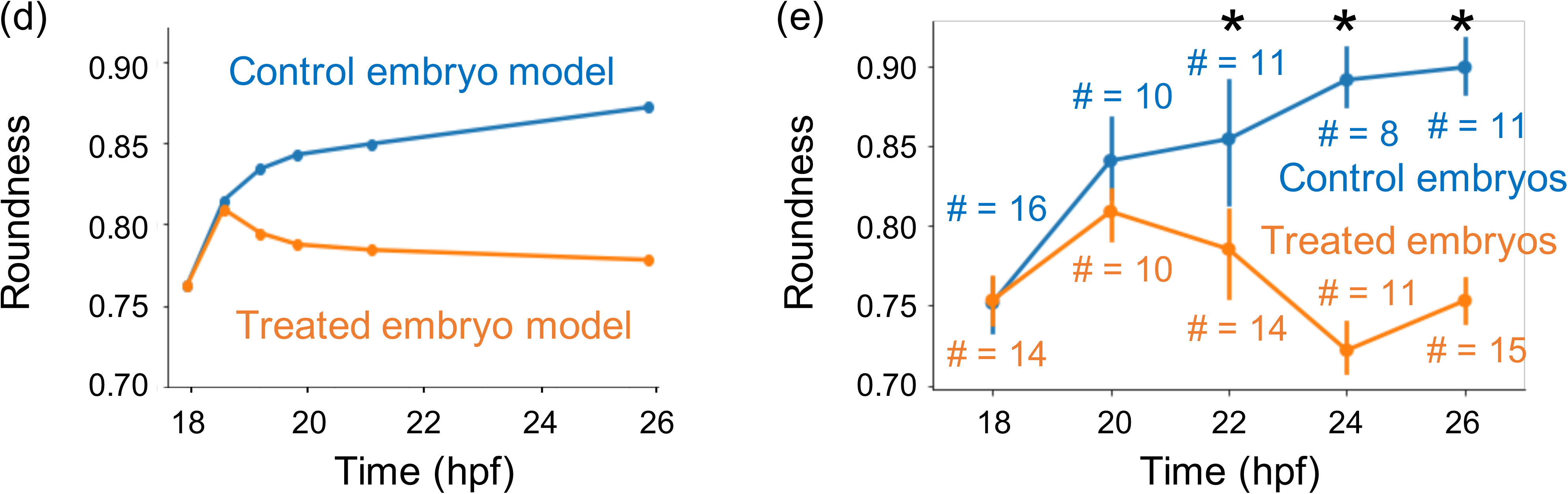
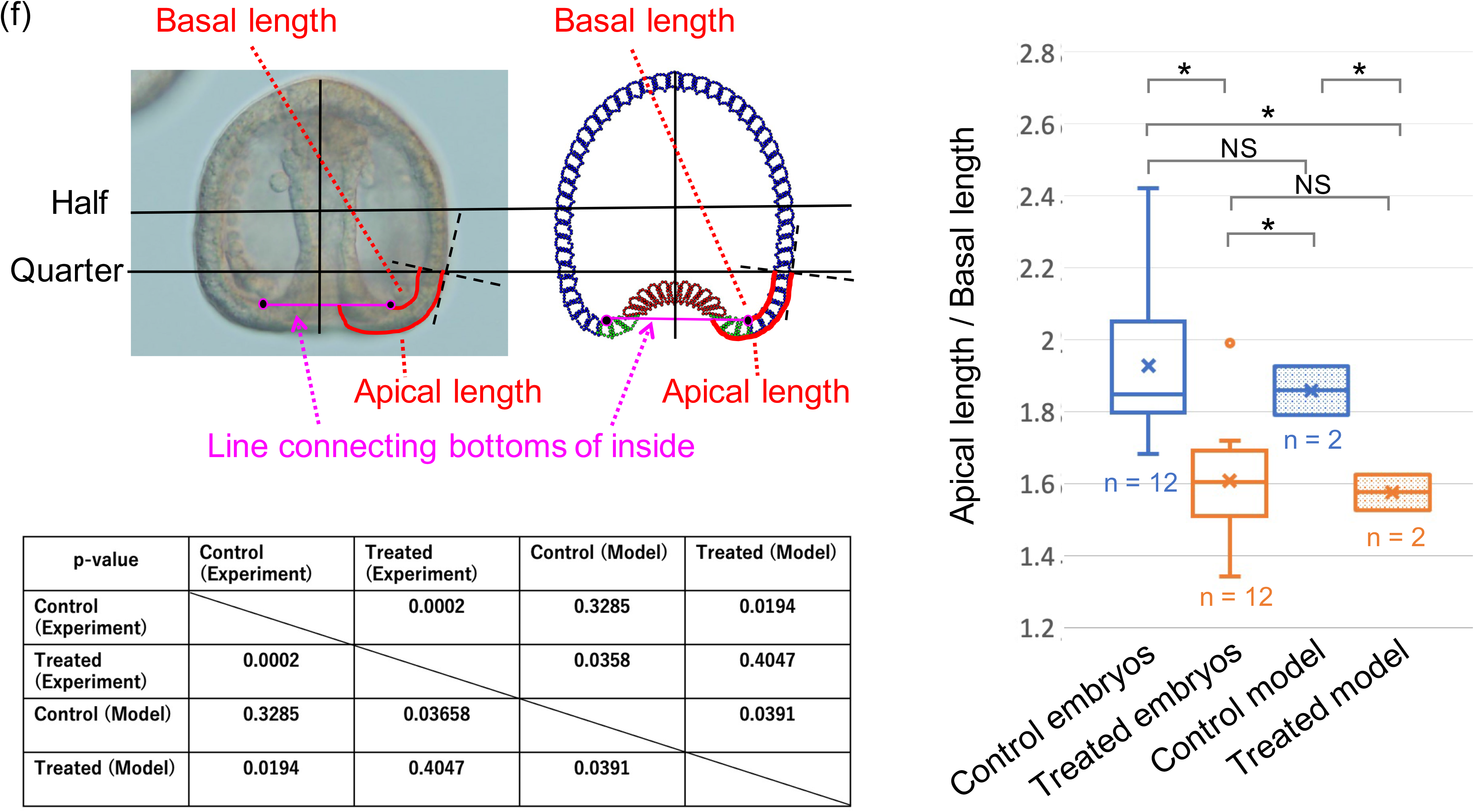
Coarse-grained model and simulation of control, treated, and over-polarized embryos. **(a)** Modeled cell lengths of apical and basal sides of control, treated, and over-polarized embryos; 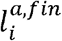 and 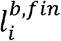 refer to the final length of the apical and basal sides of the *i*-th cell, respectively (see Materials and methods). Distributions of the final length of the former two models were determined based on the distributions of apical-basal ratios of actinin-GFP intensities from control and treated embryos in step 2. The top center panel was a modification of Fig. 3d. Red, green, and blue circles in the top panel and panel **(b)** represent pigment cells, wedge cells, and other cells, respectively. **(b)** Snapshots of the initial embryo shape at 18 and 26 hpf of the three models. **(c)** Definition of the roundness of the vegetal side of embryos from simulation results and imaging. **(d-e)** Roundness indices of vegetal sides of modeled control and treated embryos **(d)** and experimentally determined values (e). Blue and orange # refer to the sampling number of control and treated embryos at each time point, error bars indicated 95% confidence intervals, and * indicated that the roundness of control embryos was significantly larger than that of treated embryos according to Welch’s t-test (p < 0.05, see Fig. S5b). (**f**) The ratio of apical/basal lengths obtained by experimental and simulation data, and p-values of Welch’s t-test between the two types of data, where * and N. S. between two box plots indicates p < 0.05 and p > 0.05, respectively. Embryos at step 4 (late gastrula) and simulation final step were compared.

Simulations showed different shapes (Fig. 5b), although each shape was similar to that experimentally observed at step 4 (Fig. 1). Similar temporal changes in roundness indices (Fig. 5c) of control embryos (0.85 ~ 0.9) and treated embryos (0.75 ~ 0.8) were observed in the simulated (Fig. 5d, Fig. S5a) and experimental data (Fig. 5e, Fig. S5b). Additionally, the ratio of apical/basal lengths along the embryos around the vegetal pole obtained at the final time step in the simulations of both control and the treated model exhibited similar features to those of the control and treated embryos at step 4 (late gastrula), respectively (Fig. 5f). Here, the control embryos and control model showed significantly larger ratios than the treated embryos and treated model, respectively, but there were no significant differences between the observed embryos and simulation results in both the control and treated conditions.

The simulation where the elongation ratio of the apical side of each cell was a little larger than that of control was also performed, named “over-polarized embryo” (Fig. 5a, b). The whole embryonic shape of this model showed in a wider and larger roundness index than the control embryos (Fig. S5a). This suggested that the site-specific apical-basal ratio polarity of F-actin distribution in each cell influenced sensitively the whole embryonic shape.

## 3 Discussion

Omeprazole treatment resulted in abnormal embryo shape with incomplete gastrulation, in particular, more than half of treated embryos showed partial exogastrulation with outward extending endodermal tissue (Figs. 1 and 2). Such anomalous morphogenesis was due to the perturbation of intracellular F-actin distribution in the vegetal half of embryos (Figs. 3 and 5) as follows:

F-actin concentrations generating cell cortical forces were significantly larger at the apical side of each cell than that of the basal side in the vegetal half of the control embryos, except for cells at the vegetal pole. Therefore, the apical planes of wedge cells and other cells in the vegetal pole side were expected to be significantly wider than the basal planes. Conversely, the apical planes of cells in the vegetal half of the treated embryo were not significantly wider compared to the basal sides due to lower apical F-actin levels. Simulation results showed that such expected features in the vegetal half may lead to normal early gastrulation in control embryos and partial exogastrulation in treated embryos (Fig. 5). Therefore, appropriate embryonic intracellular apical F-actin distribution in the vegetal half is required for gastrulation.

Although the present mathematical model assumed that F-actin concentration correlates positively with the forces elongating each cell periphery to stretch tissues, the force by F-actin varies both qualitatively and quantitatively depending on its interaction with the environment and other molecules. At least for the morphogenetic process considered in this study, the employed assumptions for the relationship between F-actin concentration and the elongation of each side of the cells were supported by the relationships between the apical and basal lengths in the control and treated embryo after morphological changes (Fig 5f).

Gastrulation does not proceed when apical lamina is inhibited (Burke et al., 1991). The whole embryonic distributions of apical lamina along cells were nearly identical between the control and treated embryos, although large differences were observed between the cell shapes in the vegetal half and whole embryos. This indicated that the generated force was independent of the relative amounts of the apical lamina in each cell. Embryos with a partial knockout of F-actin-regulating factors (*RhoA* or *YAP1*) showed inhibited gastrulation at the primary invagination step (Fig. 4a, b). This indicated that the secondary invagination was inhibited without appropriately regulated F-actin, even if the apical lamina existed. Therefore, the apical lamina may play the role of just the scaffold for apical F-actin, and F-actin generates the force required for deformation processes driving the secondary invagination of gastrulation and whole embryonic deformation. Notably, the importance of the role of intracellular F-actin for gastrulation was also supported by the recent study performing Rab35 knockdown (Remsburg et al., 2021).

The elongation of the archenteron requires cell divisions and rearrangements (Ettensohn, 1985). Conversely, in order to focus on the morphogenetical process independent of cell divisions or those during early gastrula stage with little occurrence of cell divisions, the present mathematical model did not involve the effects of cell divisions. Therefore, the model could not reproduce cell division induced morphological processes like elongations of archenteron of normal embryos and more outward protruding of the part of the elongated archenteron in treated embryos (Fig. 1). However, the morphological features evaluated by the temporal change in the roundness index and the lengths of apical and basal sides of the vegetal side of both control and treated embryos, which should be spatially and temporally little influenced by cell divisions, were well reproduced in our models (Fig. 5d-f). These facts suggested that not only cell divisions but also the cytoskeletal forces provide dominant contributions to gastrulation. The considerations of the model implementing the effects of cell divisions are important and are considered as a future issue for not only the simulations of whole gastrulation processes but also the considerations of the regulatory mechanism of post gastrulation processes like left-right asymmetry formations of embryos as mentioned previously.

The present mathematical models were constructed based on the results of the apical-basal ratios of actinin-GFP of step 2 to simulate the morphological features of step 3 and beyond (Fig. 5a). It was noted that higher values of the apical-basal ratio of actinin-GFP intensities were obtained at step 1 than at step 2 (Fig. 3b-c). This may be explained by the fact that the cells at step 1 are larger than those at step 2 (Fig. 3b-c), because the apical-basal difference between the molecule concentrations were expected to increase with the increases in apical-basal distance. Additionally, the absolute values of actinin-GFP intensities in the apical and basal sides of cells in both control and treated embryos at step 1 were sufficiently smaller than those at step 2 (Fig. S2). These findings suggest that the contributions of the apical-basal ratios of actinin-GFP intensities at step 2 were more dominant than those at step 1 for the focused gastrulation processes.

CRISPR-Cas9-mediated *RhoA*-knockout showed an intermediate level (55.6%) of mutation frequency (Fig. S4) and the knockout embryos exhibited primary invagination. However, LV embryos expressing dominant-negative *RhoA* failed to initiate primary invagination (Beane et al., 2006). Therefore, the contribution of *RhoA* is expected to be considerable also for the primary invagination, but the extent of the dependency on F-actin function is small and a mechanism other than the F-actin-related system might be involved in this initial process, at least in HP embryos. F-actin function may be essential for secondary invagination.

F-actin network formation, generating cell cortical forces, was drastically enhanced with decreased pH *in vitro* (Köhler et al., 2012). Whole embryonic distributions of apical-basal ratios of the pH indicator at step 2 was found to correlate positively with those of intracellular actinin-GFP signal intensities (Fig. 3c, e), suggesting that F-actin concentration was high when the pH was low at the apical side of cells. Additionally, the apical-basal polarity of F-actin distribution in the vegetal half cells of the omeprazole-treated embryo decreased to significantly smaller than those of control embryo (Fig. 3c) while whole embryonic distribution of the apical lamina did not exhibit any significant changes (Fig. 3d). Conversely, the whole embryonic distribution of intracellular pH and polarity was unchanged even when F-actin polymerization was directly perturbed by the knockout of F-actin-regulating factors (Fig. 4d). These facts suggested that the intracellular pH is a one-sided regulator of F-actin polymerization also in HP embryos and contributes to the progression of the secondary invagination to form appropriate apical-basal intracellular polarity of F-actin distribution.

Herein, omeprazole increased the intracellular pH in most cells in the HP embryo (Fig. S6) conversely from the apical-basal polarities of intracellular pH. Oppositely from the case of HP embryo, a recent study of the LV embryo reported that omeprazole significantly decreases intracellular pH (Schatzberg et al., 2015). This suggests the apical-basal polarities of intracellular pH in LV embryo was increased by omeprazole. The influence of pH on F-actin network formation and force generation might be a universal biochemical feature as confirmed via both *in vitro* experiments (Köhler et al., 2012) and the present *in vivo* study. Therefore, omeprazole may increase the apical-basal polarities of intracellular pH and F-actin concentration in LV embryo similar to the “over-polarized embryo” model. The over-polarized embryo model exhibited a wider embryonic shape with a larger roundness index than the control embryos (Fig. 5b, S5b), which is consistent with LV imaging (Schatzberg et al., 2015). This also supported the significant contributions of cell position-dependent intracellular F-actin apical-basal polarizations to early gastrulation.

Omeprazole treatment of sea urchin embryos disrupts the left-right asymmetric formations of the adult rudiment although gastrulation is completed (Duboc et al., 2005; Hibino et al., 2006; Bessodes et al., 2012) (Fig. S7). The left-right asymmetric *nodal* gene expression observed immediately after gastrulation is important for establishing this asymmetry (Duboc et al., 2005). Disrupted left-right asymmetry in HP is observed at lower (60 ~ 80%) omeprazole concentrations (Hibino et al., 2006), different to what was observed in the present study. Dilute omeprazole solutions induced weak perturbation of gut formation in the present study, which may disrupt cell-cell interactions and regulate whole embryonic *nodal* gene expression.

This report revealed the cell position-dependent regulations of intracellular F-actin polymerization and polarization by intracellular pH by focusing on the normal and omeprazole-induced abnormal gastrulation in sea urchins. Frogs, chickens, zebrafish, and ascidians disrupted left-right asymmetry upon inhibition of H^+^/K^+^ ion pump activity (Levin et al., 2002; Kawakami et al., 2005; Shimeld and Levin, 2006). Therefore, omeprazole may disrupt left-right asymmetry of sea urchin embryo through the inhibition of H^+^/K^+^ ion pump activity (Duboc et al., 2005; Hibino et al., 2006; Bessodes et al., 2012), which suggests the involvement of H^+^/K^+^ ion pump in intracellular pH distribution regulating F-actin distribution. Due to the high similarity between H^+^/K^+^ ion pump and Na^+^/K^+^ ion pump sequences in sea urchin, however, the detail features of H^+^/K^+^ ion pump such as whole embryonic gene expression profiles along the developmental process and intracellular distributions have yet to be determined; hence further investigations are warranted to explore the formation mechanism underlying embryonic position-dependent intracellular states promoting gastrulation. The universal effect of inhibiting H^+^/K^+^ ion pump activity on early embryo formation among different animals should also be determined in the future.

## 4 Experimental Procedures

### 4.1 Animals and embryos

Adult Japanese sea urchins (HP) were collected from the Seto Inland Sea or Tateyama Bay. Eggs and sperms were obtained via coelomic injection of 0.55 M KCl. Fertilized eggs were cultured in filtered seawater at 16°C.

### 4.2 Omeprazole treatment

A 100 mM stock solution of omeprazole (FUJIFILM Wako Pure Chemical Corporation, Japan) dissolved in dimethyl sulfoxide (DMSO) was immediately added to fertilized eggs at a final concentration of 100 μM and cultured at 16°C.

### 4.3 Whole mount *in situ* hybridization

Whole mount *in situ* hybridization of glial cells missing (*gcm*) was performed following the method of Minokawa (2004). Antisense RNA probe of *gcm* was transcribed from a PCR fragment amplified from the cDNA clone with those primers stated in Table S1. Antisense RNA probe was synthesized using MEGAscript T7 Transcription Kit (Ambion, USA) with DIG RNA Labeling Mix (Roche, Germany).

Sea urchin embryos were fixed by fixative III (4% paraformaldehyde, 32.5% filtered seawater, 32.5 mM 3-(N-morpholino) propane sulfonic acid (MOPS) (pH 7.0), 162.5 mM NaCl) for 16 h at 4 °C. Fixed embryo were preserved in 70% ethanol at −20 °C until use. Embryos were washed three times with 100 mM MOPS, 500 mM NaCl, and 0.1% Tween20 (MOPS buffer). The embryos were prehybridized for 3 h in the hybridization buffer containing 70% formamide, 100 mM MOPS (pH 7.0), 500 mM NaCl, 0.1% Tween20, 1 mg/ml bovine serum albumin (BSA) at 50 °C. For hybridization, 0.1 ng/μl of probe was added to the prehybridized samples, and the hybridization was performed at 50 °C with shaking for 6 days. After hybridization, the embryos were washed five times with MOPS buffer. The embryos were then transferred to probe-free hybridization buffer again at 50 °C for 3 h and washed three times with MOPS buffer. For blocking, the embryos were incubated in MOPS buffer containing 10 mg/ml BSA for 20 min at RT, followed by additional incubation in the MOPS buffer containing 10% sheep serum and 1 mg/ml BSA for 30 min at 37 °C. Incubation with a 1:1,500 dilution of the alkaline phosphatase conjugated Fab fragments (Roche Molecular Biochemicals, Indianapolis, IN) in the MOPS buffer containing 1% sheep serum and 1 mg/ml BSA was performed overnight at 4 °C. Embryos were washed 5 times every 2 h to remove antibodies with MOPS buffer (the last wash was done with overnight shaking). The embryos were washed 2 times with alkaline phosphatase buffer (100 mM Tris HCl (pH 9.5), 50 mM MgCl_2_, 100 mM NaCl, and 1 mM Levamisole). The staining reaction was done in the alkaline phosphatase buffer containing 10% dimethylformamide and NBT (SIGMA, Germany) and BCIP (SIGMA, Germany).

### 4.4 Alkaline phosphatase staining

Alkaline phosphatase staining of endodermal tissue was performed following Whittaker and Meedel (1989) with modification. Sea urchin embryos were fixed by fixative III (4% paraformaldehyde, 32.5% filtered seawater, 32.5 mM 3-(N-morpholino) propane sulfonic acid (MOPS), pH 7.0, 162.5 mM NaCl) for 1 h at 4 °C. Embryos were washed three times using 0.5 mL 1x phosphate-buffered saline. Embryos were reacted in reaction buffer (100mM Tris-HCL (pH 9.5), 100mM NaCl, 5mM MgCl2) containing NBT (SIGMA, Germany) and BCIP (SIGMA, Germany). Embryos showed specific staining when reacted at 4 °C for 48h and then at RT for 6h.

### 4.5 Live imaging of pH, actinin, and apical lamina

Intracellular pH of sea urchin embryos was visualized using pHrodo Red AM intracellular pH indicator or 5-(and-6)-carboxy SNARF-1 (C-1270) (Thermo Fisher Scientific, USA) at final concentrations of 10 and 5 μM, respectively, and stained at 16 °C for 30 min. The embryos were washed in filtered seawater and observed using confocal microscopy with laser illumination at 555/585 nm excitation/emission for pHrodo and 555/573 m excitation/emission for SNARF-1, respectively.

Actinin and fibropellin-1 were visualized following the fusion of the proteins with GFP. RNA was extracted from mesenchyme blastula stage HP embryos using ISOGEN (Nippongene, Japan) according to the manufacturer’s instructions. Actinin- and fibropellin-1-coding sequences were amplified via RT-PCR using SuperScript™ III Reverse Transcriptase (Thermo Fisher Scientific, USA) with the primers stated in Table S2 and cloned into pGreenLantern2-derived plasmid at EcoRI and XhoI restriction sites. mRNA (actinin-GFP, fibropellin-1-GFP, in that order) was synthesized *in vitro* using an mMESSAGE mMACHINE T7 ultra transcription kit (Thermo Fisher Scientific, USA) and purified using an RNeasy mini kit (Qiagen, Nederland). mRNA was microinjected into fertilized eggs as described by Liu et al. (2019).

Fluorescent images were acquired by confocal microscopy using an excitation of 488 nm, and emission of 515 nm. The images of the animal and vegetal poles of the embryo were analyzed based on the *z*-stacked image with the largest area by averaging seven *z*-axis images at a total thickness of 6 μm.

### 4.6 Quantification of fluorescent signals

Fluorescent images of embryos were acquired according to the angle θ (0°-180°) between the vegetal and animal poles along the circumference (Fig. 3a, Fig. S7) and transformed into a band-like image using the “Polar Transformer” function (https://imagej.nih.gov/ij/plugins/polar-transformer.html) of ImageJ 2.1.0. Each band-like image was filtered using a median filter (radius = 1.0). The filtered images were binarized using “Mean” (pHrodo indicator), “IsoData” (actinin), and “Triangle” (fibropellin-1 and SNARF-1) functions, respectively, to obtain the cellular regions of the embryo. The apical-basal ratios of the pH indicator, actin-GFP, and fibropellin-1-GFP for θ were determined using average fluorescent intensities over the regions at a width of 3.126 μm from the apical and basal sides (Fig. 3a, Fig. S7).

The fluorescent intensity of the pHrodo indicator was defined as the average fluorescent intensity over the intracellular region at a width of 6.232 μm from the apical side (Fig. S7). Fluorescent signal intensity values were estimated by calculating the ratio between the observed fluorescence intensity and the average background fluorescence intensity around the entire embryo.

F-actin and apical lamina were stained via mRNA microinjections of actinin-GFP and fibropellin-1-GFP into fertilized eggs. Nonnegligible variations was inevitable in the concentration of injected mRNA among fertilized egg samples, which would be amplified during development. Therefore, only the apical-basal ratios were mainly used to evaluate the intracellular features of F-actin and apical lamina distributions. For the fluorescence intensity of actinin-GFP in the apical and basal sides of cells, see Fig. S2.

Two data points were obtained for each θ from each fluorescence image of the embryo since the left and right sides of the embryo were considered axis symmetric against the animal-vegetal axis. Both data were used to estimate the sample average and 95% confidence interval of each value, where the number of samples (n) was given as 2 × [number of observed embryos].

### 4.7 Gene knockout using CRISPR-Cas9

Knock out of endogenous *RhoA* and *YAP1* in HP was performed using the method described by Liu et al. (2019). The oligonucleotide sequences used for sgRNA preparations are listed in Table S3.

### 4.8 Heteroduplex mobility assay (HMA) and DNA sequencing

Analysis of knockout embryos using HMA and DNA sequencing was performed as previously published by Liu et al. (2019). The primer sequences used to amplify each target region are listed in Table S4.

### 4.9 Mathematical model of sea urchin embryos

A two-dimensional particle model describing cellular motion at the cross-section of animal poles and vegetal poles of sea urchin embryos during steps 1 to 3 of gastrulation was constructed. The determined number of cells was based on that at the equatorial plane of the embryo at the blastula stage (Mizoguchi, 1999). The following assumptions were made: 1) each cell was represented by 16 particles with radii (r) of 1.125 μm, and each embryo was represented by 64 cells connected in a ring (Fig. S8); 2) each cell perimeter was 36 μm and the height and width of the embryo at step 1 were 110 and 100 μm, respectively, which was consistent with sea urchin embryo observations (results not shown); 3) the number of cells was constant because cell divisions and cell invasions from other cross-sections were rarely observed during the primary invagination stage (Mizoguchi, 1999); 4) the motion of each particle obeyed the overdamped limit of the equation (1) of motion,

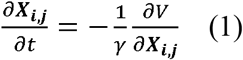

where ***X***_*i,j*_ = (*x*_*i,j*_(*t*), *y*_*i,j*_(*t*)) is the position of the *j*-th particle constructing the *i*-th cell (*i* = 0,1,2…63 and *j* = 0,1,2…15) on the x-y plane at time *t* (Fig. 5, Fig. S8), *γ* is the coefficient of the drag force acting on each particle, and V is the potential of the entire system; 5) the y-axis was parallel to the animal-vegetal axis of the embryo model.

We rewrote 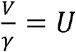 and *U* was calculated as follows:

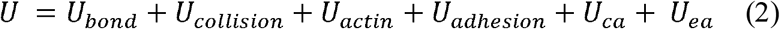

where *U*_*bond*_ is the elastic force potential between each neighboring pair of particles in each cell to maintain each cell perimeter, denoted as follows:

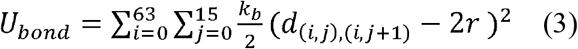

where *k*_*b*_, *d*_(*i,j*),(*i,j*+1)_, and *r* are the coefficient of elasticity, the distance between the *j*-th and *j*+1-th particle (J+ 1 = 0 for J= 15) in the *i*-th cell at time *t*, and the particle radius, respectively.

*U*_*collision*_ is the potential of excluded volume effects among all particles denoted as follows:

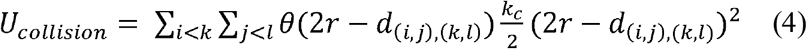

where *k*_*c*_ is the coefficient of repulsion between two particles, and *θ*(*x*) is the Heaviside step function.

*U*_*actin*_ is the elastic force potential to form and sustain cell shape with the expansion and contraction of the apical and basal sides by the cytoskeleton, denoted as follows:

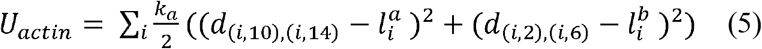

where the apical and basal sides of the *i*-th cell consisted of particles with *j* = 10 –14 and *j* = 2 – 6, respectively. The wideness of the apical and basal sides at time *t* were given by *d*_(*i*,10),(*i*,14)_ and *d*_(*i*,2),(*i*,6)_, respectively. The basic wideness of the apical and basal sides were given by 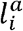 and 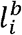, respectively. *k*_*a*_ was assumed by the coefficient of elasticity to sustain the the wideness of apical and basal sides of the cell.

*U*_*adhesion*_ is the potential force for cell adhesion by proteins, such as cadherin, denoted as follows:

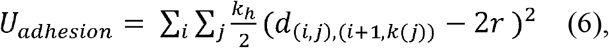

where *k*_*h*_ is the coefficient for the adhesive forces between the particles of the *i*-th and *i* + 1-th cells (*i* + 1 = 0 for *i* — 63), and *k*(*j*) ≡ 6, 7, 8, 9, 10 for *j* = 2, 1, 0, 15, 14, respectively (Fig. S8).

*U*_*ca*_ is the potential of the forces to conserve each cell area, denoted as follows:

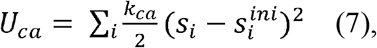

where *k*_*ca*_ is the coefficient of elasticity required to maintain each cell area. The area of the *i*-th cell was estimated by 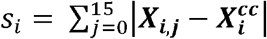 with 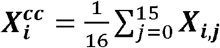, and 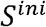 referred to *S*_*i*_ given by ***X***_*i,j*_ at the initial state.

*U*_*ea*_ is the potential of the forces to maintain the area (volume) of the sea urchin embryo, denoted as follows:

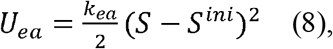

where *k*_*ea*_ is the coefficient of elasticity required to maintain embryo area. The area was estimated by 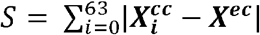 with 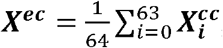, and 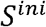 referred to *S* given by ***X***_*i,j*_ at the initial state.

*U*_*ca*_ and *U*_*ea*_ were considered alternatives for the volume-conserving forces in real 3-D cells and embryos as internal pressures in each cell and embryo were isotropic.

### 4.10 Simulation method for the mathematical model

The simulations of the present mathematical model were performed through the integrals of the equation of motion (1) using the Euler method at time intervals of 0.000064 h with conserved ***X***^*ec*^ = (0, 0). In all models, the parameters *k*_*bond*_, *k*_*collision*_, *k*_*actin*_, *k*_*adhesion*_, *k*_*ca*_, and *k*_*ea*_ were given empirically as 9375, 6250, 8125, 8125, 625, and 0.00625 hour^−1^, respectively, because there were no experiments to measure or estimate them. The qualitative features of the results were independent of the details of these values if the order was maintained. The model formed the embryo shape at step 1 of gastrulation if the appropriate cell type-dependent values of 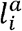 and 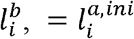 and 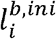 were given for the models of pigment cells, wedge cells, and other cells (Fig. 5). This configuration gave the particle positions ***X***_*i,j*_ at time t = 0 (initial configuration) in all simulations.

The early gastrulation processes were simulated by the change in and 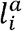 and 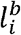 respectively, from = 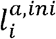 and 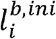 to = 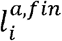 and 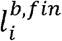 at time *t* = 0. Here, 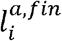 and 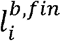 were assumed to obey 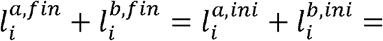 constant among cells except pigment cells based on the expectation that F-actin was constant. The model showed similar structural behaviors to the early gastrulation as the relaxation process of 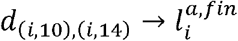 and 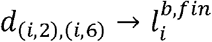.

### 4.11 Statistical analysis

All experiments were performed independently twice or more with four replicates or more per experiment. Statistical test was performed using SciPy library (https://www.scipy.org/).

The roundness index of the vegetal side of each embryo was evaluated using the following: [embryo width half-way between the bottom and middle of the embryo]/[embryo length] (Fig. 5c).

The apical length of embryos around vegetal pole was defined as the length of the curve along apical side from 1/4 of the height of the whole embryo to a point that this curve crosses with straight line connecting the bottom of the inner side of the embryo (Fig. 5f). The basal length was defined as the curve from the bottom of the inner side of the embryo to the point at which this curve intersects the line that crosses the tangent line of the apical envelope at 1/4 of the embryo perpendicularly (Fig. 5f). The same definitions were employed to determine the apical and basal lengths around vegetal pole also for simulation results (Fig. 5f). Each length of the curve was measured using ImageJ 2.1.0. The number of samples (n) was provided as 2 × [number of observed embryos], and those from the simulation results was considered 2 since the data were obtained from left side and right side of each model.

## Acknowledgments

We thank Prof. Masato Kiyomoto (Tateyama Marine Laboratory, Ochanomizu University) for supplying live sea urchins. We would like to thank Editage (https://www.editage.com) for English language editing.

## Author Contributions

K.W., N.S., and A.A. conceived and designed the study; K.W., Y.K., and N.S. conducted the experiments; K.W., Y.Y., M.F., and A.A. analyzed the data; K.W. and A.A. conducted the mathematical model construction and simulations; K.W., N.S., and A.A. wrote the manuscript with support from all authors; T.Y. supervised the work.

## Data Availability

The datasets generated during and/or analyzed during the current study are available from the corresponding author on reasonable request.

