## Supplementary information for "Partial exogastrulation due to apical-basal polarity of F-actin distribution disruption in sea urchin embryo by omeprazole"

Supplementary Material

### Supplementary Figures and Tables

#### Supplementary Figures

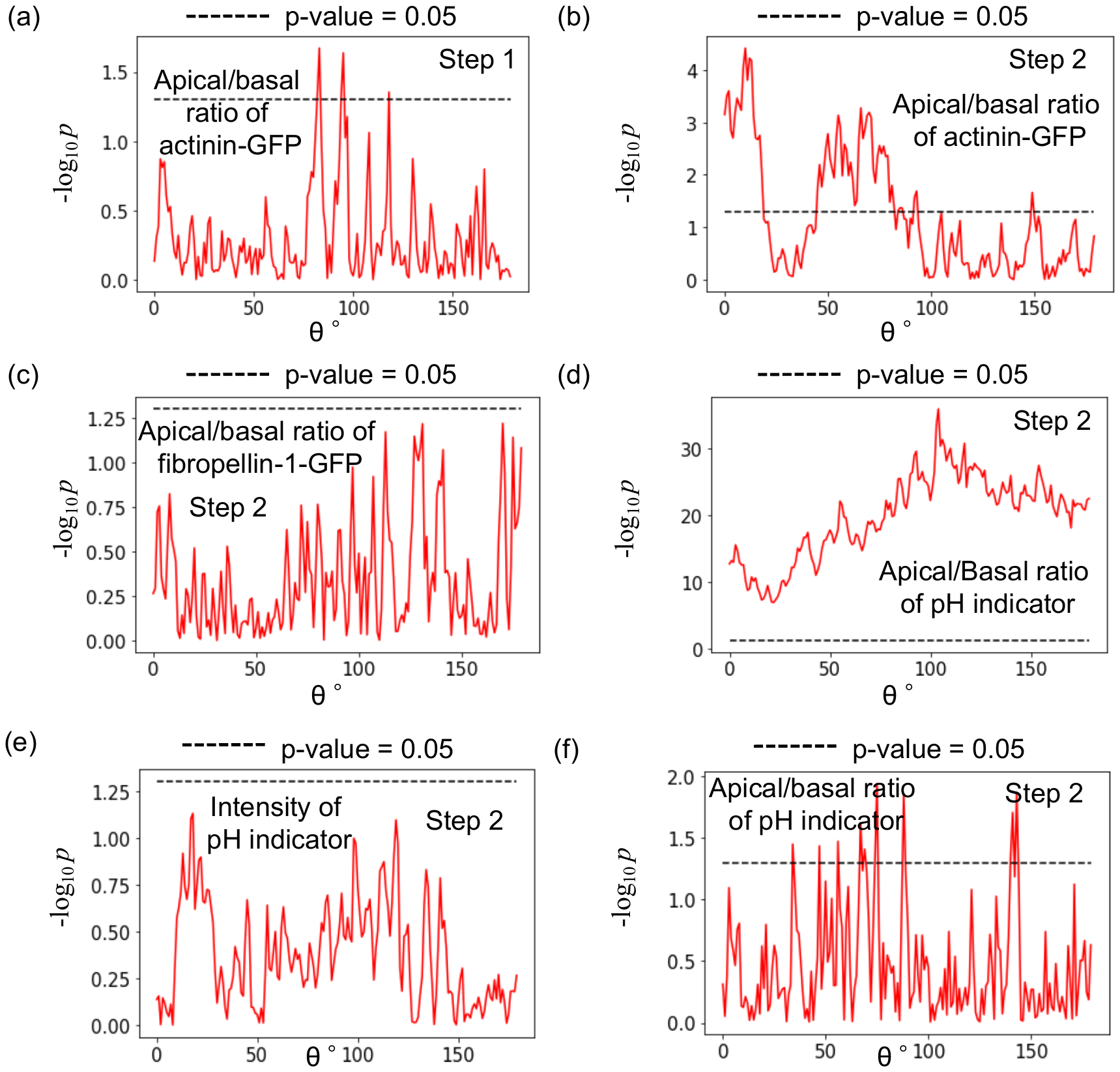

**Fig. S1. Distributions of p-values (-log_10_ *p*).** **(a-d)** P-value distributions (-log_10_ *p*) of apical/basal ratios obtained using Welch’s t-test for control and treated embryos at each *θ* (0°-180°), comparing fluorescence intensities of actinin-GFP of steps 1 **(a)** and 2 **(b)**, fibropellin-1-GFP **(c)**, and pH indicator **(d)**. **(e-f)** Distributions of p-values (-log_10_ *p*) of Welch’s *t*-test of *RhoA* knockout embryos and control embryos at each *θ* comparing fluorescence intensities of the pH indicator **(e)** and the apical/basal ratio of the pH indicator **(f)**.

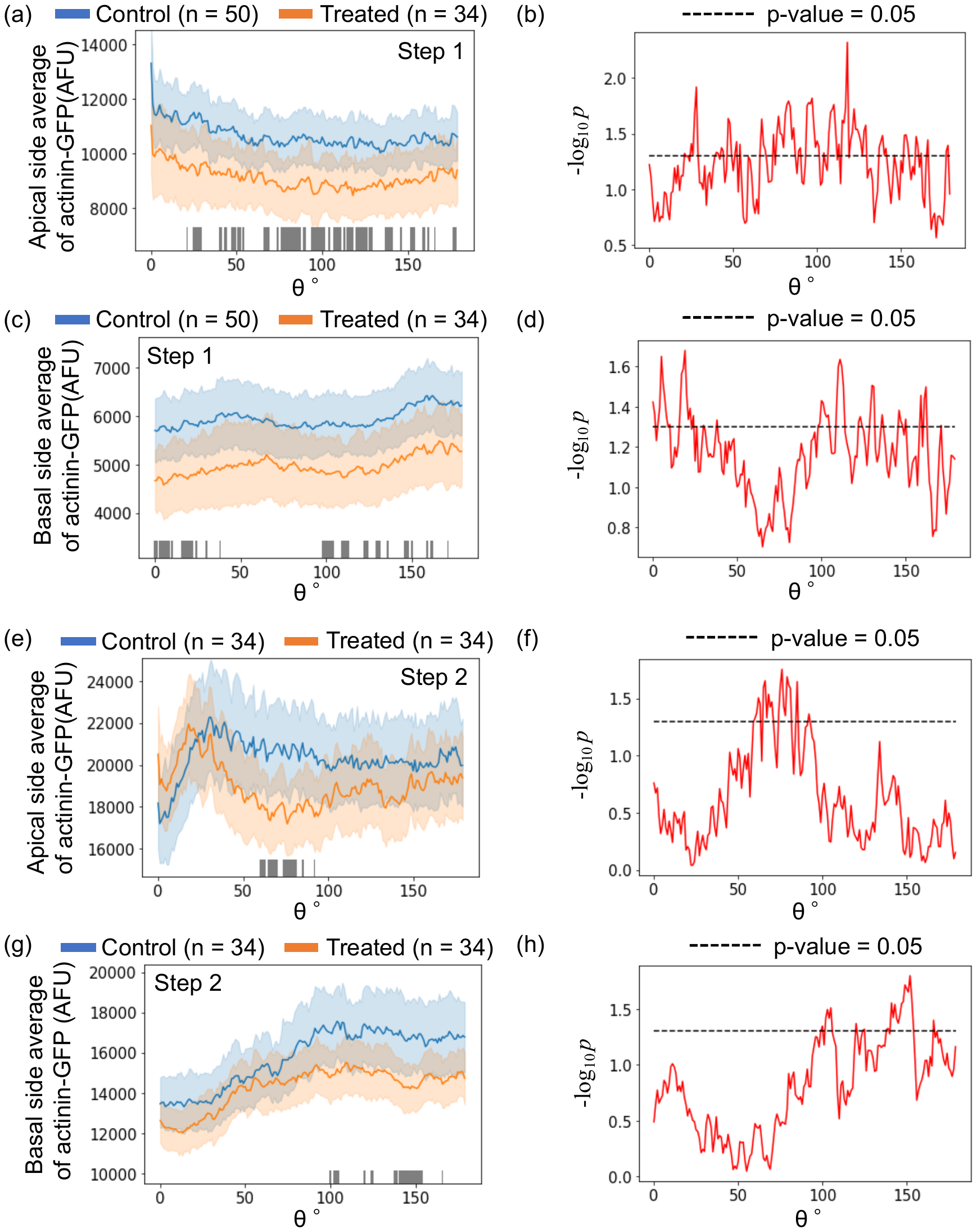

**Fig. S2. Evaluation of intracellular actinin-GFP intensities.** Distributions of intracellular fluorescence intensities of actinin-GFP in control and treated embryos in apical side at step 1 **(a)**, in basal side at step 1 **(c)**, in apical side at step 2 **(e)**, and in basal side at step 2 **(g)** at each *θ***.** P-value distributions (-log_10_ *p*) of the intracellular actinin-GFP intensities in apical side at step 1 **(b)**, in basal side at step 1 **(d)**, in apical side at steps 2 **(f)**, and in basal side at step 2 **(h)** at each *θ* obtained using Welch’s t-test. The indications of the colors and *θ* are stated in Fig. 3.

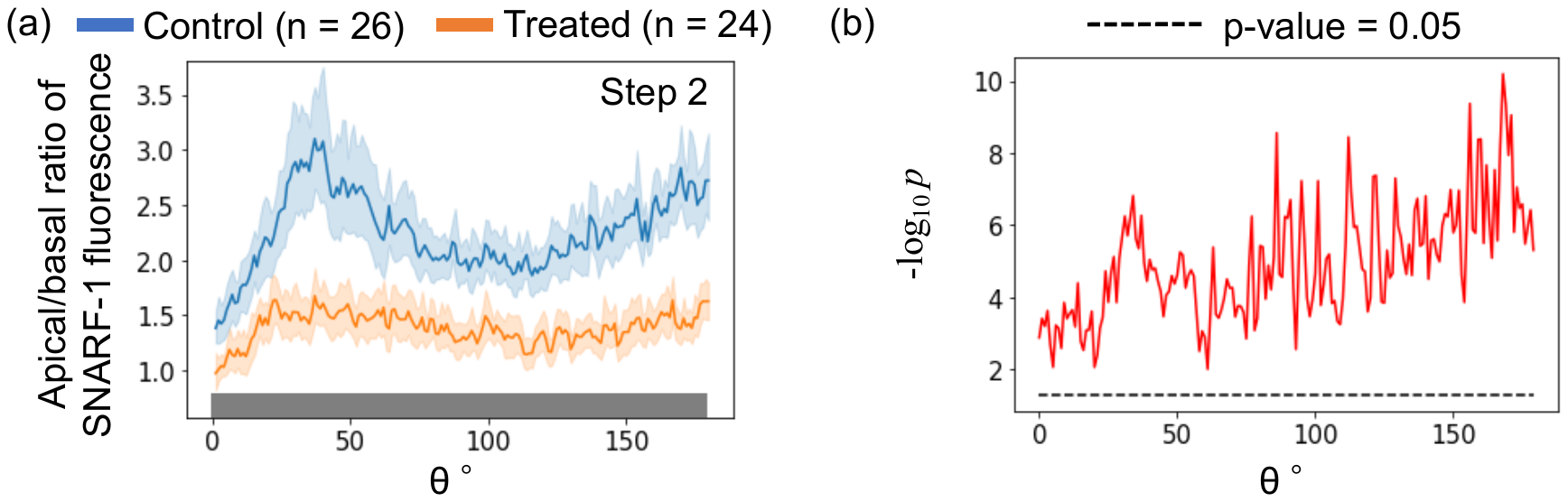

**Fig. S3. Evaluation of intracellular pH polarity via SNARF-1.** **(a)** Distributions of intracellular polarities of SNARF-1 fluorescence intensity as a function of angle *θ* in control and treated embryos at step 2 where the pH was higher at lower SNARF-1 intensity. **(b)** P-value distributions (-log_10_ *p*) obtained using Welch’s *t*-test between control and treated embryos for intracellular polarities of SNARF-1 intensity at each *θ*. The indications of the colors and *θ* are stated in Fig. 3. The correlation coefficients between the apical/basal ratio of SNARF-1 and actinin-GFP signal intensities (Fig. 3E) in control and treated embryos were 0.48 and 0.64, respectively.

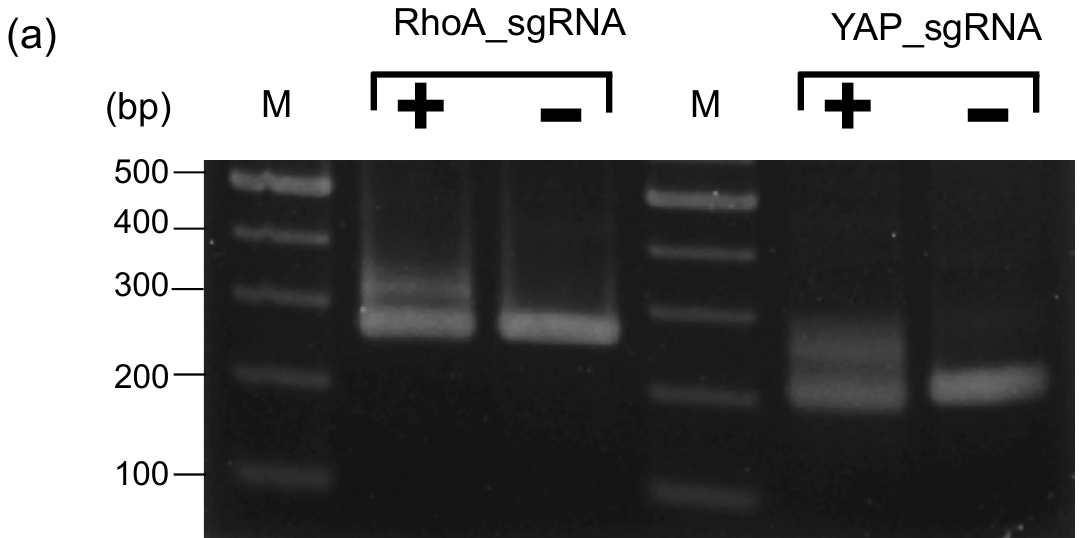

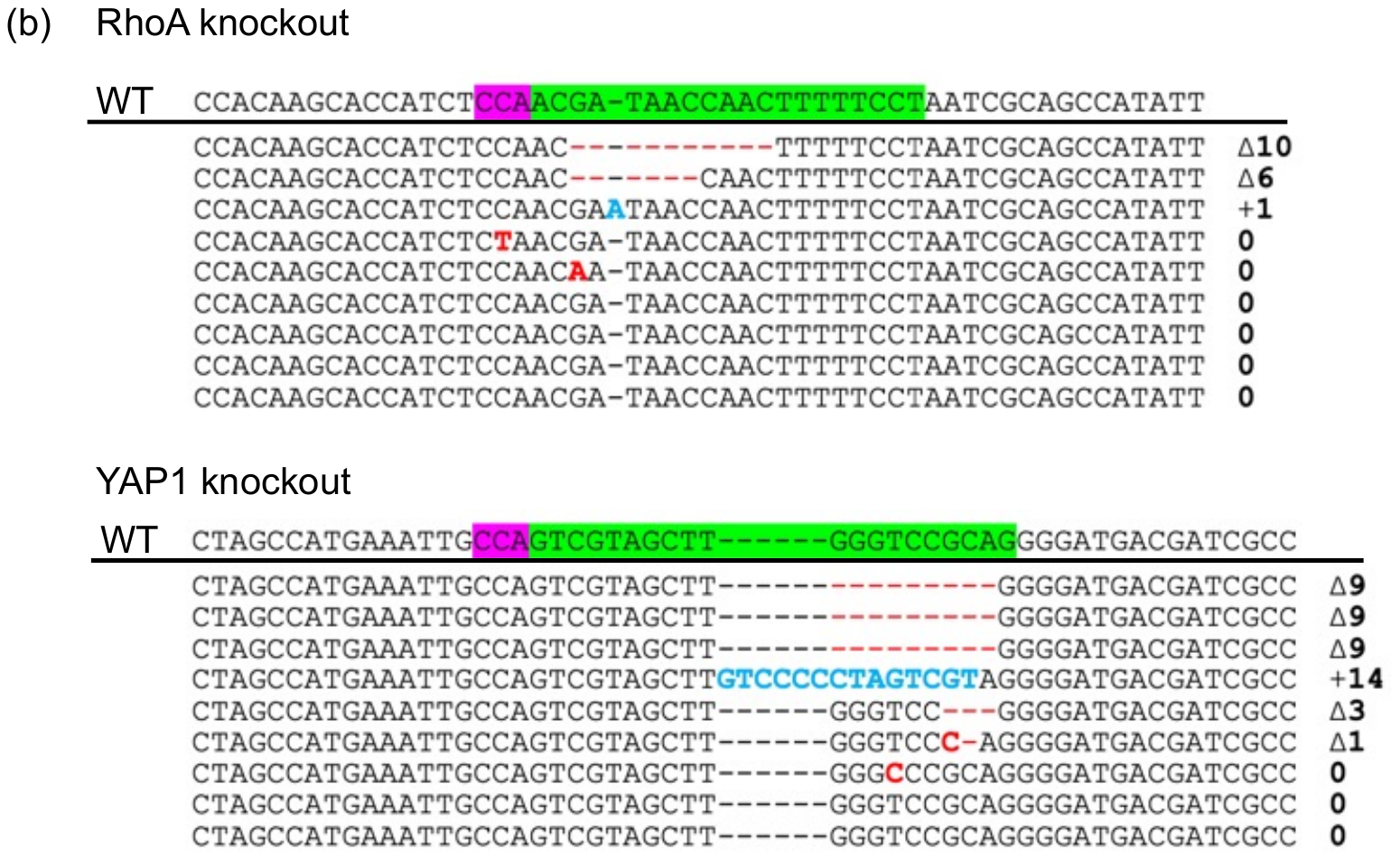

**Fig. S4. Mutation efficiency in knockout embryos.** **(a)** Genotyping using heteroduplex mobility assay (HMA) of genomic DNA extracted from individual sea urchin embryos injected with Cas9 alone at 40 hpf (control, -) or Cas9/sgRNA (*RhoA* or *YAP1* knockout, +) using the primers listed in Table S3. PCR products were separated on a 3% agarose gel. M indicates 100-bp DNA ladder. **(b)** Sequences of 9 *RhoA* knockout and 9 *YAP1* knockout embryos. The wild-type (WT) sequences are shown on top with the protospacer adjacent motif (PAM) highlighted in magenta and the protospacer highlighted in green. Deletions, substitutions, and insertions are indicated by red dashes, red letters, and blue letters, respectively.

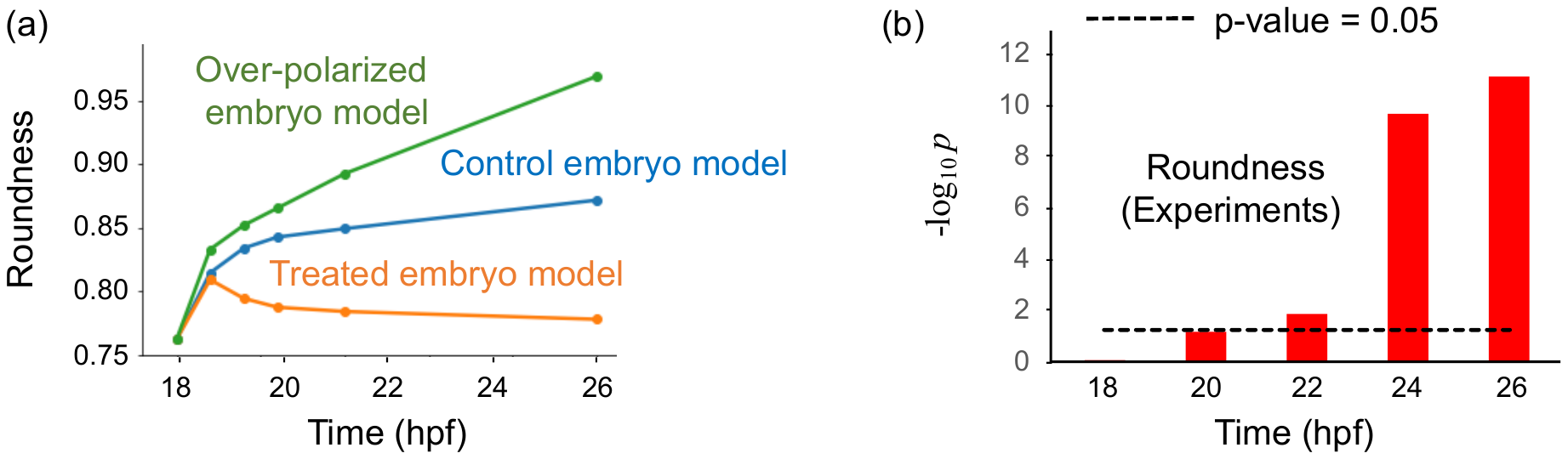

**Fig. S5. Simulation of the roundness index and p-values of the vegetal side of embryos.** **(a)** Time course of modelled vegetal side roundness of control, treated, and over-polarized embryos. **(b)** P-values (-log_10_ *p*) of roundness indices between vegetal sides of control and treated embryos determined using Welch’s *t*-test.

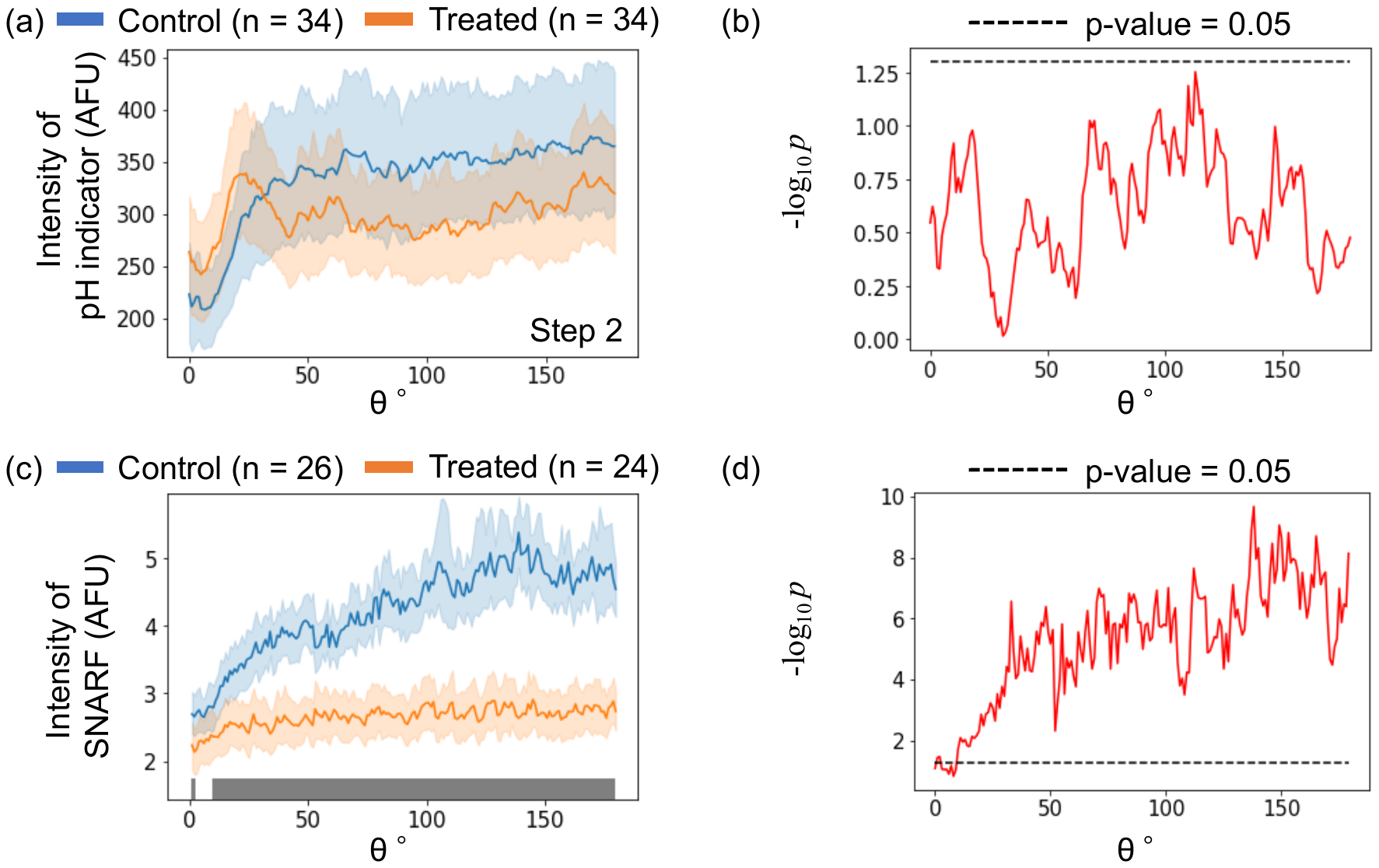

**Fig. S6. Evaluation of intracellular pH.** Distributions of intracellular fluorescence intensity of pH indicator **(a)** and SNARF-1 **(c)** in control and treated embryos at step 2 at each *θ*. P-value distributions (-log_10_ *p*) of the intracellular intensity of pH indicator **(b)** and SNARF-1 **(d)** between control and treated embryos at each *θ* obtained using Welch’s t-test. The indications of the colors and *θ* are stated in Fig. 3.

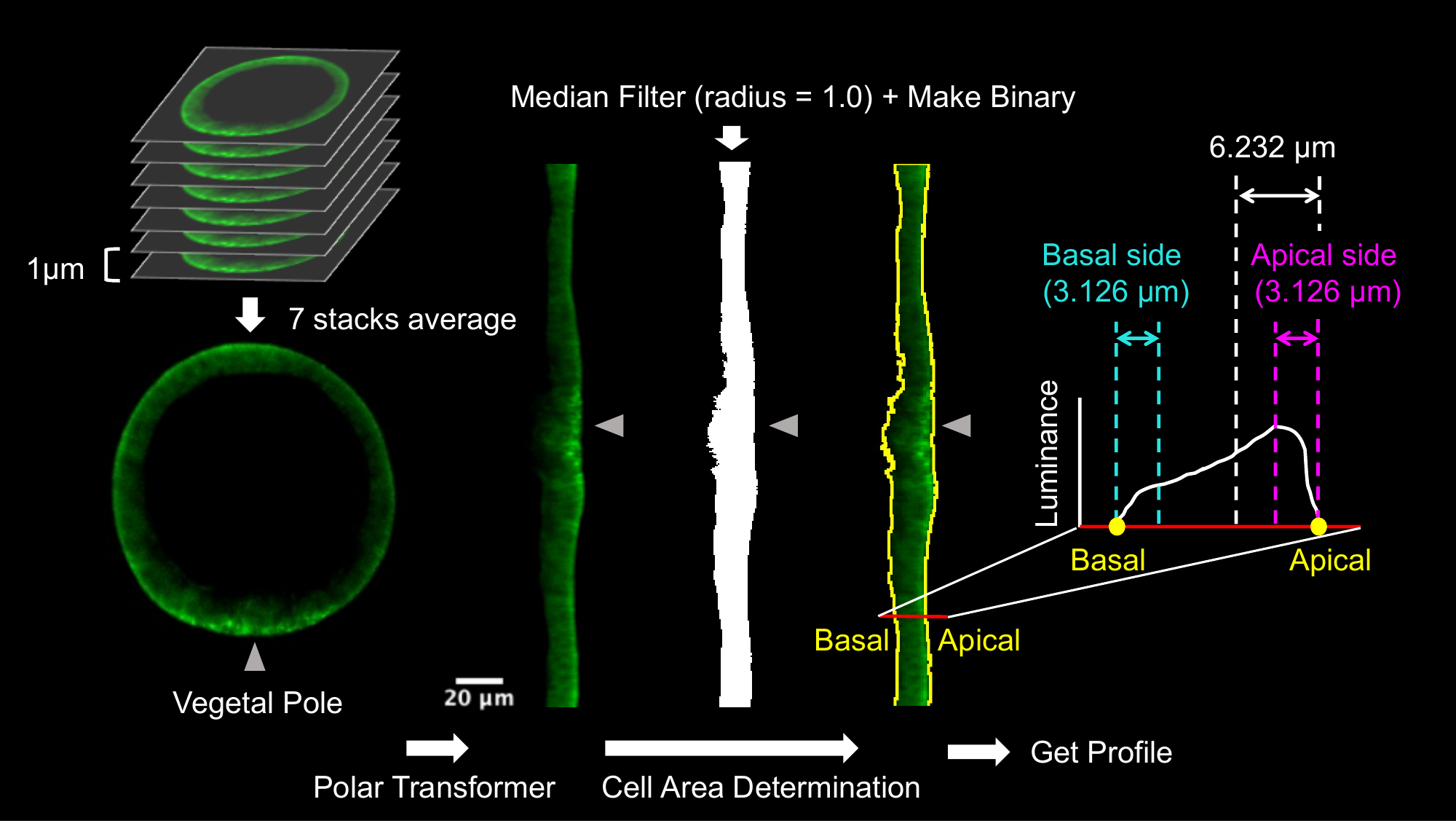

**Fig. S7. Quantification of fluorescent images.** Workflow of the image analysis method used to evaluate intracellular fluorescence intensities and polarities (see Materials and Methods).

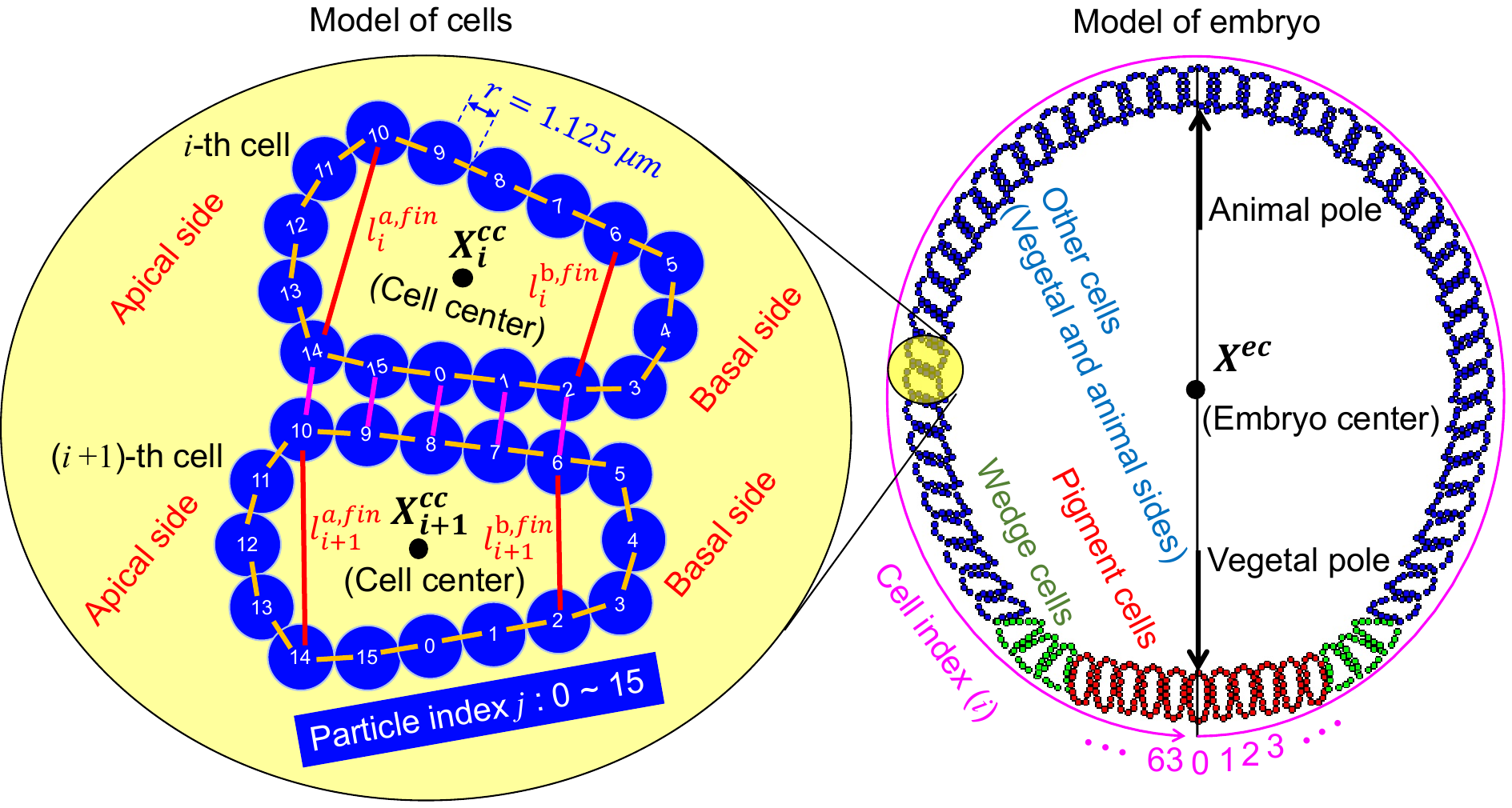

**Fig. S8. Illustration of mathematical model geometry** **for each cell and embryo consisting of 64 cells.** Each pair of particles joined by orange or magenta lines was connected by a spring with a natural length equal to their diameters. Each loop formed by 16 particles along orange lines represent the membrane of each cell, and the elastic force of the spring between the pair of particles connected by the magenta line describes the adhesive force between the cells. The cell cortical force at the apical and basal sides of each cell was modeled by the elastic force of the spring between each pair of particles connected by a red line with natural length ${=l}_{i}^{a,fin}$ and $l_{i}^{b,fin}$, respectively. The black point marked “$\boldsymbol{X}_{\boldsymbol{i}}^{\boldsymbol{cc}}$**”** represents the position of the center of the *i*-th cell. The magenta curve around the model of the embryo starting from the vegetal pole is the axis of the cell index *i* = 0, 1, … 63. The black point marked “$\boldsymbol{X}^{\boldsymbol{ec}}$**”** represents the center of the embryo. Red, green, and blue cells represent pigment, wedge, and other cells, respectively, which show the different mechanical features during gastrulation.

**Table S1. Primers used in the PCR assembly of the antisense RNA probe**

| Name | Forward primer sequence (5’ to 3’) | Reverse primer sequence (5’ to 3’) |
| --- | --- | --- |
| GCM | CGACTGATAACCACGCTCAAC | TCACCATCTATCCACTCGTT |

**Table S2. Primers used for cDNA isolation**

| Name | Forward primer sequence (5’ to 3’) | Reverse primer sequence (5’ to 3’) |
| --- | --- | --- |
| Actinin-GFP | CTAGGTACCAAGATCGCCACCATGGCGTACTATGGCAATC | GCCGCCACTAGAATTGCAAGCTCACTCTGGCCG |
| Fibropellin-1-GFP | CTAGGTACCAAGATCGCCACCATGAGGACGTGGTTACTAGC | GCCGCCACTAGAATTGCTGCATCAGGCTGAGGTG |

**Table S3. Oligonucleotides used in the PCR assembly of the sgRNA template**

| Name | Nucleotide sequence (5’ to 3’) |
| --- | --- |
| RhoA knockout | GTAATACGACTCACTATAGGGAAAAAGTTGGTTATCGTGTTTTAGAGCTAGAAATAG |
| YAP knockout | GTAATACGACTCACTATAGGCGTAGCTTGGGTCCGCAGGTTTTAGAGCTAGAAATAG |

**Table S4. Primers used in HMA**

| Name | Forward primer sequence (5’ to 3’) | Reverse primer sequence (5’ to 3’) |
| --- | --- | --- |
| RhoA_HMA | GGGCATGTGTATTTCCTACAAACGGC | CCTGTTTACCGTCTACTTCTATATCAGC |
| YAP_HMA | AATAAGCCATTCGAGGGAGTCGAGCGCAG | TCGTAGCTCTGTTGACGAAGATGCTGATT |
